# Testicular mRNA-LNP Delivery: A Novel Therapy for Genetic Spermatogenic Disorders

**DOI:** 10.1101/2025.05.21.654986

**Authors:** Chenwang Zhang, Nan Liang, Wenbo Li, Shuai Xu, Peng Li, Wanze Ni, Na Li, Sha Han, Ningjing Ou, Haowei Bai, Yuxiang Zhang, Furong Bai, Yifan Sun, Dewei Qian, Xinjie Bu, Erlei Zhi, Ruhui Tian, Yuhua Huang, Jingpeng Zhao, Fujun Zhao, Hao Chen, Zheng Li, Chencheng Yao

## Abstract

Uniform testicular maturation arrest is a severe form of male infertility characterized by the presence of germ cells that do not complete spermatogenic development. It is usually caused by meiotic arrest with genetic variants and difficult to treat via drugs or surgery. mRNA-lipid nanoparticle (LNP) delivery is a promising therapeutic option for maturation arrest with monogenic variants via protein replacement therapy. Herein, a spermatocytes-tropic LNP (Pool1-LNP3) was identified via a library of 30 ionizable lipids screening. And *in vivo* delivery of this novel LNP composition using rete testis microinjection was showed to be high spermatocytes targeting with high transfection efficiency. Thereafter, it was revealed that *in vivo* delivery of Pool1-LNP3 encapsulating *Msh5* mRNA could promote crossover formation and restore spermatogenesis in *Msh5^D486Y/D486Y^* mouse models with DSB recombination defects. Notably, the offspring without genomic integration was born using intracytoplasmic sperm injection (ICSI) derived from rescue of *Msh5^D486Y/D486Y^*mouse and embryo transfer. Furthermore, no obvious inflammation and histologic damage in any tissue were detected after *in vivo* delivery of mRNA-LNP. In addition, it was demonstrated that *Maps* mRNA-LNP3 recovered spermatogenesis in *Maps* KO mouse with meiotic arrest. Altogether, these findings suggested that this spermatocytes-tropic mRNA-LNP delivery could become a viable and broad applicable strategy for treatment of spermatogenic disorders with genetic defects, providing a foundation for future clinical application.

## Introduction

Infertility constitutes a profound global health burden, affecting 10-15% couples worldwide and contributing to significant psychological distress and socioeconomic impacts^1^. Approximately 50% of infertility-affected cases involved male factors^2^. And spermatogenic disorders is assumed to be the most severe type. Spermatogenesis is an intricate process orchestrated by over 2,000 genes. Spermatogenic disorders could result in non-obstructive azoospermia (NOA), which is classified as Sertoli cell only syndrome (SCOS), hypo-spermatogenesis (HS), and mature arrest (MA) according to testis biopsy and subsequent pathologic analysis. Uniform testicular maturation arrest was a subset of NOA, and it was susceptible to genetic defects, including microdeletion of Y chromosome, chromosomal translocation, and monogenic disorder^3,4^. The application of whole-exome sequencing (WES) technology has led to the identification of numerous pathogenic variants in essential regulators of spermatogenesis, for instance: MutS Homolog 5 (*MSH5*), Chromosome 3 Open Reading Frame 62 (*C3ORF62*), Meiotic Nuclear Divisions 1 (*MND1*), Meiosis Inhibitor Protein 1 (*MEI1*), and Synaptonemal Complex Protein 2 (*SYCP2*) ^5–10^. These genetic variants are usually associated with defects of the events of meiosis, including synapsis complex formation and DSB combination, which leaded into meiotic arrest and NOA^11^. Current interventions, including hormone therapy such as aromatase inhibitors and microdissection testicular sperm extraction (micro-TESE) achieve limited success in genetically driven uniform testicular maturation arrest^12–14^. Given the vast heterogeneity of genetic variants implicated in spermatogenic disorders and the current lack of broad-spectrum therapeutic strategies tailored to distinct genetic etiologies in clinical practice, there is an urgent need to develop a broad applicable treatment modality capable of addressing diverse genetic variants underlying spermatogenic disorders.

Gene therapies primarily address the genetic defects of disease pathogenesis rather than merely alleviating symptoms. Previous study revealed that CRISPR/Cas9 combined with *in vivo* electroporation could be used to restore spermatogenesis by correcting the mutant locus in *Msh5^D486Y/D486Y^*mouse model based on pathogenic variants identified in patients with spermatogenic disorder^8^. However, the clinical application of germline gene editing remains constrained by off-target risks and ethical prohibitions in genome editing of human gametes and zygotes. Consequently, developing alternative therapeutic strategies for diverse pathogenic variants is imperative. Protein replacement therapy (PRT) emerges as a promising approach for treating spermatogenic disorders with genetic defects. mRNA-based delivery is the widely applicated PRT utilizing carriers to deliver protein-coding mRNA into target cells for transient translation. Following cellular delivery, mRNA is directly translated into protein in the cytoplasm, where each protein undergoes post-translational modifications and native folding *in vivo* to perform its physiological functions accurately. This intrinsic capacity to generate fully functional proteins *in situ* underscores a critical advantage of mRNA-based therapeutics over exogenous protein supplementation^15^. Crucially, mRNA therapeutics enable transient protein expression without genomic integration, minimizing mutagenic risks associated with permanent genetic alterations^16,17^. Recent advancements in chemical modifications (e.g., pseudouridine incorporation) and lipid nanoparticle (LNP) delivery systems have remarkedly improved mRNA stability, translational efficiency, and cell-type specificity^18^. Thus, mRNA-based PRT could be used as a particularly suitable strategy for rescuing spermatogenic disorders caused by genetic variants^19^.

Microinjection into seminiferous tubules of naked mRNA followed by electroporation has showed partial efficacy in rescue of spermiogenesis defects in *Armc2*-knockout mouse models^20^. However, this approach induced localized testicular necrosis and exhibited suboptimal delivery efficiency, raising substantial concerns regarding safety and reproducibility in human applications. These findings underscore the critical need for advanced delivery platforms to overcome the inherent instability and low efficacy associated with unformulated mRNA therapies. Adeno-associated virus (AAV) vectors, while efficient in gene delivery, raise concerns over immunogenicity, risk of genomic integration and limited cargo size^21–24^. However, lipid nanoparticles validated by their clinical success in COVID-19 vaccines, have revolutionized nucleic acid delivery by enabling efficient mRNA encapsulation, endosomal escape, and controlled protein expression^25^. Recent advances in mRNA-LNP have expanded their utility beyond liver-centric diseases such as lungs, placenta and brain ^26–28^. However, in the context of treating spermatogenic failure caused by genetic variants, research on mRNA-LNP delivery remained largely unexplored. Although previous studies have reported that LNP-mediated delivery of self-amplifying RNA (saRNA) successfully rescued the meiotic arrest phenotype in *DNA meiotic recombinase 1* (*Dmc1*) knockout (KO) mice, several concerns remain^29^. It has yet to be verified whether the prolonged overexpression of saRNA within germ cells might lead to toxicity, and *in vivo* safety of the virus-derived nonstructural proteins encoded by the saRNA has not been fully established. Moreover, given that the saRNA contains alphavirus replicase genes and encodes an RNA-dependent RNA polymerase (RdRP) complex with a length approaching 7 kb, delivering larger target gene sequences would undoubtedly pose a significant challenge to this approach.

Herein, we reported the use of a *in vivo* screening approach to develop an mRNA-LNP platform to restore spermatogenesis in *Msh5^D486Y/D486Y^* mouse models derived from NOA-affected patients^8^. Patients and mice carrying variant in *MSH5* presented meiotic arrest phenotype and are infertile. Using a rationally designed LNP library, we identified formulations with enhanced tropism for spermatocytes, achieving transient restoration of functional protein expression without genomic integration. *In vivo* delivery of Pool1-LNP3 encapsulating *Msh5* mRNA could promote crossover formation and restore spermatogenesis in *Msh5^D486Y/D486Y^*mouse models with DSB recombination defects. Notably, the offspring without genomic integration was born using intracytoplasmic sperm injection (ICSI) derived from rescue of *Msh5^D486Y/D486Y^* mouse and embryo transfer. Furthermore, no obvious inflammation and histologic damage in any tissue were detected after *in vivo* delivery of LNP. In addition, it was demonstrated that *Maps* mRNA-LNP3 recovered spermatogenesis in *Maps* KO mouse with meiotic arrest. Therefore, this approach not only circumvents the limitations of viral vectors but also establishes a framework for repeatable, broad-spectrum treatment of genetic male infertility. Our findings highlighted the potential of mRNA-LNPs to overcome therapeutic barriers in clinical reproductive medicine, offering a paradigm shift in addressing spermatogenic disorders with genetic defects.

## Materials and methods

### Mouse model and cell line

All animal care and experiments complied with the guidelines of the National Institutes of Health and were approved by the Animal Care Committee of Shanghai General Hospital (2022AWS0287). The *Maps* KO mouse was constructed based on the CRISPR/Cas9 technology, following the knockout strategy described in previous study^30^. DNA isolated from tail were used for genotyping by PCR and Sanger sequencing as described previously^8,30^, the primers listed in Supplementary Table 1. All mice were maintained under controlled environmental conditions (temperature: 24 ± 1 □; relative humidity: 50-60%) with a standard 12-hour light-and-dark cycle. The HEK-293T, TM3 and TM4 cells were cultured with Dulbecco’s Modified Eagle Medium (DMEM, Gibco) supplemented with 10% fetal bovine serum (FBS, HyClone) under standard culture conditions (37 □, 5% CO_2_).

### Lipid nanoparticle synthesis

Lipid nanoparticles (LNPs) were prepared by mixing lipids in an ethanol phase with mRNA in an aqueous phase in two syringe pumps. In brief, the ethanol phase was prepared by solubilizing a mixture of ionizable lipid, cholesterol (AVT Pharmaceutical Tech), 1,2-distearoyl-sn-glycero-3-phosphocholine (DSPC, AVT Pharmaceutical Tech), and 1,2-dimyristoyl-rac-glycero-3-methoxypolyethylene glycol-2000 (DMG-PEG 2000, AVT Pharmaceutical Tech) at a predetermined molar ratio in ethanol (50: 38.5:10:1.5). The aqueous phase was prepared in 10 mM citrate buffer (pH 4) with enhanced green fluorescent protein mRNA (*EGFP* mRNA, CATUG Biotechnology), *Msh5*, and *Maps* mRNA. Syringe pumps were used to mix the aqueous and ethanol phases at a ratio of 3:1. The resulting LNPs were dialyzed against 20 mM Tris in a 100,000 MWCO dialysis tube at 4 °C for 4 h, and refreshed in 20 mM Tris buffer, following dialysis at 4 °C overnight. A RiboGreen RNA assay (Invitrogen) was used to calculate the nucleic acid encapsulation, and a dynamic light scattering (DLS, Zetasizer) was used to measure the size and polydispersity index (PDI) of the LNPs (Supplementary Table 2).

### *In vivo* luciferase mRNA LNP delivery

Pool1-LNP3 encapsulating luciferase mRNA were administered by rete testis microinjection. 4 μg luciferase mRNA-LNP 3 was injected into left testis of each wildtype mouse. On day 1, 3, 5, 7 and 9 after injection, fluorescence imaging was captured using living image software on In Vivo Imaging System (IVIS, PerkinElmer). Ten minutes before imaging, d-luciferin potassium salt (Thermo Fisher Scientific) was injected intraperitoneal (i.p.) to mice at a dose of 150 mg/kg.

### RNA extraction and mRNA production

Total RNA was isolated from mouse testicular tissue using TRIzol reagent (Invitrogen) following the manufacturer’s protocol. RNA integrity and purity were verified using a NanoDrop (Thermo Fisher Scientific). Reverse transcription was performed using 1 μg RNA with a First Strand cDNA Synthesis Kit (Thermo Fisher Scientific).

For mouse *Msh5* and *Maps* mRNA production, the double-stranded DNA in vitro transcription (IVT) template was synthesized and cloned into a modified pUC19 plasmid vector containing a T7 RNA polymerase promoter sequence. The plasmids were linearized by BspQI (New England BioLabs) and purified using Monarch PCR & DNA Cleanup spin columns (New England BioLabs). IVT was performed using a HiScribe T7 mRNA Kit with CleanCap Reagent AG (New England BioLabs), and UTP was all replaced by N1-methyl-pseudouridine. RNA was purified using Monarch RNA Cleanup spin columns (New England BioLabs). mRNA product integrity was validated using native agarose gel electrophoresis and the mRNA was stored frozen at −80 □ for later use.

### DNA ploidy assay

DNA ploidy assay was performed as previously described with some modifications^31^. In brief, after the removal of tunica albuginea, testicular tissues were enzymatically digested with 1 mg/mL collagenase type IV (diluted in DMEM) at 37 °C for 5 min. Following centrifugation for 1 minutes at 100 g, the supernatant was removed. Subsequently, 0.6 mg/mL trypsin was added to digest the tubules at 37 °C for 10 min, and FBS was added for stopping digestion. The suspension was filtered through a 40 μm cell strainer and centrifuged for 10 min at 500 g. Finally, cells were suspended in 1× phosphatic buffer solution (PBS) and labeled with propidium iodide (PI) by 1 μg/1 × 10^5^ cells for 5 min on ice. FACS measurements were performed on BD FACSMelody^TM^ Cell Sorter. Gate was set as previously reported^31^.

### *In vivo* rete testis microinjection and preparation of histologic sections

Briefly, all mice were anesthetized via i.p. administration of avertin (250 mg/kg) and positioned in dorsal recumbency. Following aseptic preparation, a single incision was made approximately 1.5 cm up to the genitals. The testes were pulled out by holding the fat pad. mRNA-LNP or PBS was injected into the rete testis using a glass capillary under a stereomicroscope. The harvested testes were fixed in 4% paraformaldehyde (PFA) for 24 hours at 4°C, paraffin-embedded, and sectioned at 5 μm thickness using a rotary microtome (RM2255, Leica). Major organs, including the heart, liver, spleen, lungs, kidney and brain of wildtype mice were collected 24 h post injection and major organs of *Msh5^D486Y/D486Y^* and *Maps* KO mice were collected on day 21 post injection. All the tissues were fixed in PFA for 24 h and paraffin-embedded, 5 μm sections were generated using a rotary microtome (RM2255, Leica).

### Immunofluorescence staining

Before staining, tissue sections were dewaxed in xylene, rehydrated using a gradient series of ethanol solutions, and washed in distilled water. After rehydration, sections were processed for antigen retrieval with 10% sodium citrate (pH 6.0) at 115□°C for 15□min. Non-specific binding was blocked with 5% normal donkey serum containing 0.3% Triton X-100 for 2 hours at room temperature. Primary antibodies (Supplementary Table 3) were applied overnight at 4 °C, followed by species-matched Alexa Fluor-conjugated secondary antibodies (1:1000, Invitrogen) for 2 hours. Nuclear counterstaining utilized hoechst 33342 (1:1000, Roche), with imaging performed on a Leica SP8 confocal system equipped with LAS X software.

### Hematoxylin and eosin staining

Tissue sections (including testis, epididymis, heart, liver, spleen, lungs, kidneys, and brain) were dewaxed in xylene and rehydrated through a graded ethanol series followed by distilled water rinses. Sections were immersed in hematoxylin solution for 6 minutes, briefly differentiated in acid alcohol, and blued in tap water. Subsequently, slides were counterstained in acidified eosin solution for 45 seconds, dehydrated through ascending ethanol gradients, cleared in xylene, and mounted. Whole-slide digital images were acquired using a Panoramic SCAN slide scanner (3D Histech). Histopathological evaluation for tissues lesions was performed by the experienced pathologists.

### Meiotic chromosomal spread analysis

The tunica albuginea of the mouse testes was removed using microsurgical forceps, then the testes were transferred into 6-cm dish and washed by PBS. All the tissues were meticulously isolated into single seminiferous tubule using microsurgical forceps and subsequently suspended in hypotonic buffer (30 mM Tris–HCl, 50 mM sucrose, 17 mM trisodium citrate dehydrate, 5 mM EDTA, 2.5 mM dithiothreitol and 1 mM phenylmethylsulfonyl fluoride) for 25 min. The seminiferous tubules were teared into small pieces and pipetted 15 μL cell suspension spread on a glass slide in a thin layer of PFA solution containing 0.15% Triton X-100. Slides were placed in a humid chamber and kept for at least 3 h at room temperature. After air dry, the slides were stored at −80 °C for further experiments. The slide was then blocked with 5% Bovine Serum Albumin (BSA) and incubated with primary antibodies (Supplementary Table 3) overnight at 4 °C. After washing, the cells were incubated with the secondary antibody for 1 hour at room temperature. Images were captured with Leica SP8 confocal system.

### Elisa

Serum samples of wildtype mice were collected 24 h after *EGFP* mRNA-LNP3 delivery. Mouse IL-1 beta ELISA Kit (Servicebio) was used to evaluate the IL-1β level in the blood serum following the instructions of the manufacturer.

### Western blot

Testes and other major organs (heart, liver, spleen, lungs, kidney and brain) were collected from wildtype mice 24 h post-injection of PBS, 2 μg *EGFP* mRNA-LNP 3 and 4 μg *EGFP* mRNA-LNP 3, biological replicates of each group were 3. Tissues were lysed with RIPA Lysis and Extraction Buffer (Thermo Fisher Scientific) with 1% protease inhibitor mixture (Thermo Fisher Scientific) in accordance with the manufacturer’s guidelines. Protein extracts were acquired from the supernatants and were boiled for 10 min, and equal amounts from each sample were loaded onto 10% SDS–PAGE gels for gel electrophoresis (80 V in the stacking gel and 120 V in the resolving gel). After electrophoresis, the proteins in the gels were transferred to polyvinylidene difluoride membranes (Bio-Rad) and blocked using a 5% milk solution for 1 h at room temperature. These membranes were then incubated with primary antibodies (Supplementary Table 4) at 4 °C overnight and subsequently with secondary antibodies according to the protocol provided by the manufacturer. β-actin was used as the loading control.

### Intracytoplasmic sperm injection (ICSI) and embryo transfer

Female B6D2F1 mice were super-ovulated by the injection of 5 IU equine chorionic gonadotropin (San-Sheng Pharmaceutical) and followed by 5 IU human chorionic gonadotropin (hCG, San-Sheng Pharmaceutical) 48□h later. Cumulus-oocyte complexes were collected from oviducts at 14–16□h after hCG injection, and they were placed in HEPES-buffered CZB medium and treated with 0.1% hyaluronidase (Sigma-Aldrich) to disperse cumulus cells. Sperm were collected from the testis of *Msh5^D486Y/D486Y^* male mice (11 weeks old) at 3 weeks after mRNA-LNP injection and cultured in HEPES-buffered CZB medium for 15 min. The sperm head was separated from the tail by the application of several Piezo pulses, and the head was then injected into the oocyte according to the method described by Kimura and Yanagimachi^32^. Following micromanipulation, all injected oocytes were cultured in G1-plus (Vitrolife) under standard culture conditions. Pronucleus formation was checked at 6 h after ICSI, and outcomes were scored up to the blastocyst stage. For embryo transfer, two-cell embryos were surgically transferred into the oviduct of each 8 weeks old pseudo-pregnant female.

### Genotyping of single sperm and single blastocyst

Genotyping of single sperm and single blastocyst was performed using a Multiple Annealing and Looping-Based Amplification Cycles (MALBAC) WGA kit (XK-028-24, Yikon) following the manufacturer’s protocol. Briefly, individual sperm or blastocyst were lysed in a 5 μL reaction mixture containing 4.5 μL lysis buffer and 0.5 μL lysis enzyme. The samples were incubated at 50 °C for 20 min, followed by enzyme inactivation at 80 °C for 10 min. To avoid sample loss, cell suspensions were not vortexed after transfer and were briefly centrifuged. For amplification, 60 μL of reaction mix (60 μL processing buffer and 2 μL processing enzyme per sample) was added to the lysate. Thermal cycling was performed under the following conditions: initial denaturation at 94 °C for 3 min; 8 cycles of 20 sec at 10 °C, 30 sec at 30 °C, 40 sec at 50 °C, 2 min at 70 °C, and 20 sec at 95 °C; followed by 17–21 cycles (17 cycles for single blastocysts, 19–21 cycles for single sperm cells) of 20 sec at 94 °C, 15 sec at 58 °C, and 2 min at 72 °C; and a final extension at 72 °C for 5 min. Cycle numbers were optimized based on starting material quantity, as recommended for low-input samples. Amplified products were stored at 4 °C until downstream genotyping analysis.

For genotyping: thermal cycling was performed under the following conditions: initial denaturation at 94 °C for 3 min; 35 cycles of 30 sec at 94 °C, 30 sec at 60 °C, 30 sec at 72 °C and a final extension at 72 °C for 5 min. Genotyping strategy was described in the method ‘mice models’ as described above.

### Statistical analyses

Statistical analysis was performed using GraphPad Prism v.9.0. A two-tailed Student’s t-test or a one-way analysis of variance (ANOVA) was performed when comparing two groups or more than two groups respectively and data were represented mean ± standard deviation n ≥3 biologically independent mice per group. Significance thresholds: \**P* < 0.05, ** *P* < 0.01, *** *P* < 0.001, **** *P* < 0.0001.

## Results

### Identification of *in vivo* spermatocyte tropic LNP

Currently, testicular mRNA LNP delivery lacks efficient, safe, and germ cell-specific vehicles. To address these challenges, we constructed a combinatorial LNP library comprising 30 distinct formulations with different ionizable lipid architectures. Each LNP formulation was systematically prepared with four essential components: ionizable lipid, cholesterol, DSPC, and DMG-PEG-2000 with 50: 38.5: 10: 1.5 molar ratio, encapsulating enhanced green fluorescent protein (EGFP)-encoding mRNA (Fig 1a). A comprehensive analysis was performed to characterize physicochemical properties of the LNP, including size (nm) and polydispersity index (PDI) (Fig 1b).

**Fig 1.**
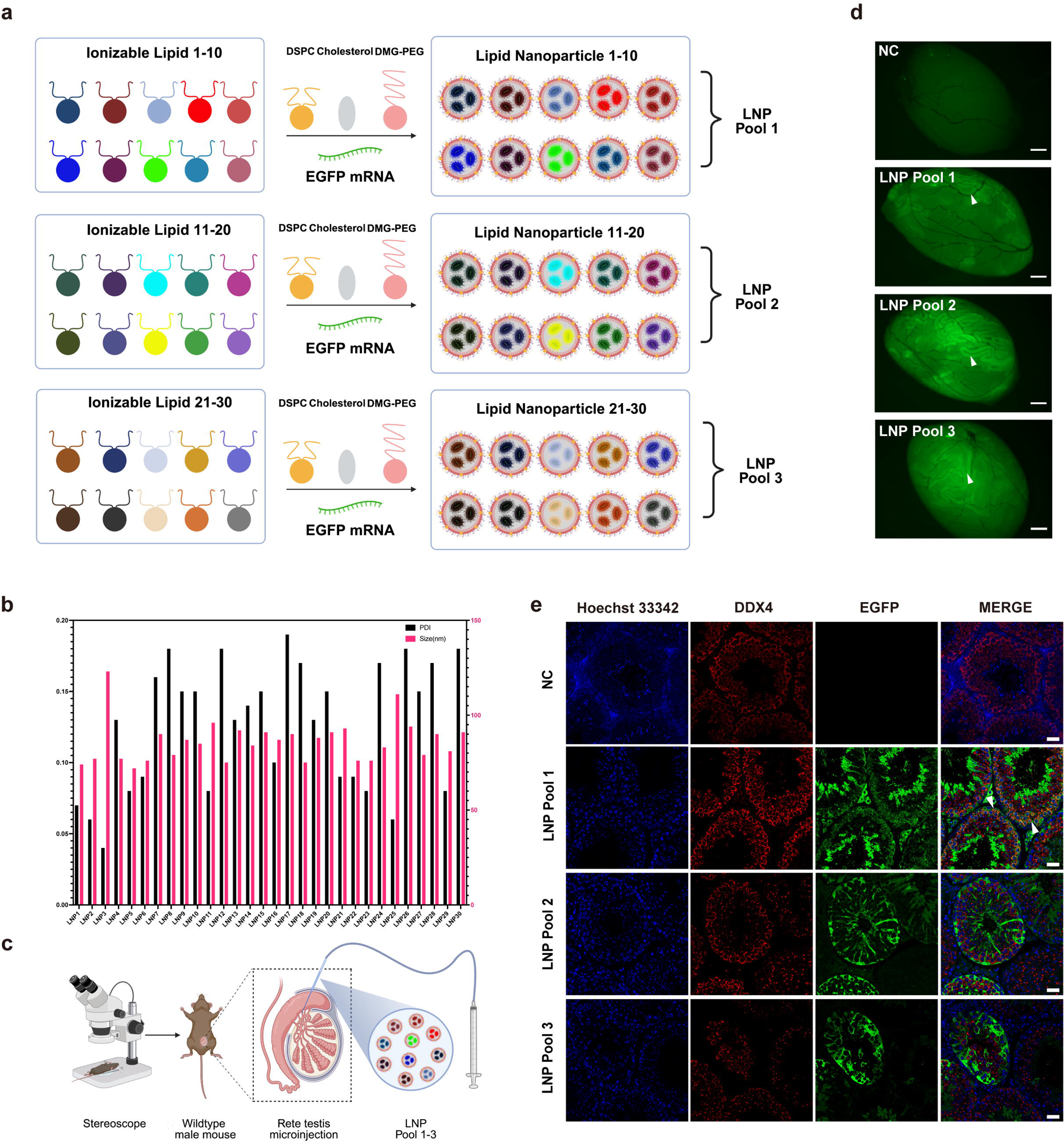
*In vivo* screening to identify a spermatocyte-tropic LNP formulation for the restoration of spermatogenic disorders. **(a)** *In vivo* screening was employed to identify a spermatocyte-tropic formulation LNP from a library of total 30 distinct LNP formulations. 30 LNP formulations were categorized into 3 LNP pools (Pool 1–3), each comprising an equimolar combination of 10 LNPs, (Created by Biorender.com). **(b)** Size and polydispersity index (PDI) of 30 LNP formulations. **(c)** Schematic diagram illustrating rete testis microinjection into seminiferous tubules of the testis, (Created by Biorender.com). **(d)** EGFP expression was observed in the seminiferous tubules 24 hours after injection of 3 LNP pools, NC denotes the negative control group administered with PBS (each group n = 3), white arrowheads indicate the EGFP positive seminiferous tubules, scale bar = 5 mm. **(e)** Immunofluorescence staining of EGFP and the germ cell marker DEAD-Box Helicase 4 (DDX4) in testicular sections of 3 LNP pools, white arrowheads indicated the EGFP expressionin spermatocytes of Pool 1, NC denoted the negative control group administered with PBS (each group n = 3), scale bar = 20 μm.

All 30 LNPs were divided into three combinatorial pools (Pool 1-3) and transfected into three kinds of cell lines (human embryonic kidney cell line (HEK293T), mouse Sertoli cell line (TM4) and mouse Leydig cell line (TM3) to confirm their expression of EGFP (Supplementary Fig 1a-c). EGFP was expressed in 3 cell lines after delivery of three LNP pools, but with different efficiency (Supplementary Fig 1d-f). To evaluate the delivery efficiency *in vivo*, LNP of three pools were separately injected to the seminiferous tubules of adult wildtype male mice via rete testis microinjection (Fig 1c). Mice were euthanized after 24 hours injection, and testicular wholemount fluorescence imaging confirmed EGFP expression across seminiferous tubules in all three LNP pools (Fig 1d). To delineate the cellular specificity of LNP-mediated mRNA delivery, testicular tissues underwent immunofluorescence staining with germ cell specific markers DEAD-Box Helicase 4 (DDX4), which was highly expressed in the cytoplasm of spermatocytes^33,34^. The results showed that LNP of Pool1 emerged as the only candidate presenting preferential mRNA delivery to germ cells (Fig 1e). To further identify the spermatocyte tropic LNP, we injected all 10 LNPs of Pool1 individually to the seminiferous tubules of wildtype male mice. Immunofluorescence staining confirmed that Pool1-LNP3 (hereafter referred to as LNP3) exclusively targeted spermatocytes (Fig 2a, b and Supplementary Fig 2e), however, EGFP was mainly expressed in Sertoli cells after injection of other LNPs of Pool1 (Supplementary Fig 2b-d). To further assess the duration of intracellular mRNA expression following mRNA delivery, we encapsulated luciferase-encoding mRNA within LNP3 and injected it into the seminiferous tubules via rete testis microinjection. Images were recorded on day 1, 3, 5, 7 and 9 after injection by *in vivo* imaging system (Fig 2c). It was showed sustained luciferase activity from day 1 to day 7 after injection (Fig 2d, e). Altogether, these results demonstrated that LNP3 were superior vehicles for mRNA delivery into spermatocytes and the sustained expression could be effective for the following rescue studies.

**Fig 2.**
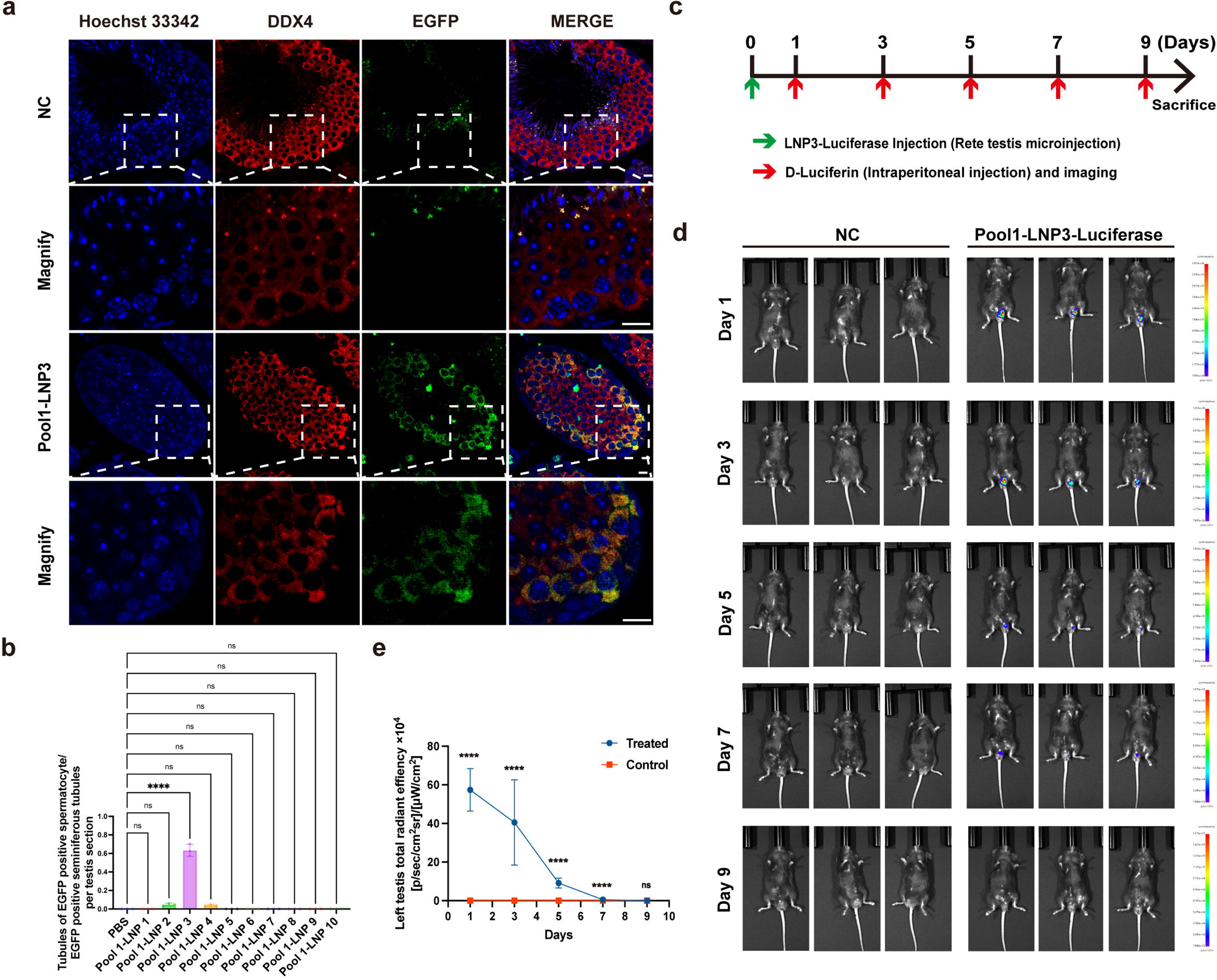
Timeline of biodistribution and expression of mRNA-LNP3. **(a)** Immunofluorescence staining of EGFP expression in testis sections after delivery of *EGFP* mRNA LNP 3, NC denoted the negative control group administered with PBS (each group n = 3), scale bar = 20 μm. **(b)** The percentage of seminiferous tubules containing EGFP-positive spermatocytes relative to the EGFP-positive seminiferous tubule population, data are represented as mean ± standard deviation, n = 3 biologically independent mice per group, one-way ANOVA and Tukey’s multiple comparisons test. ****p < 0.0001. **(c)** Schematic diagram of 4 μg firefly luciferase mRNA-LNP3 delivery via rete testis microinjection, imaging were performed followed by intraperitoneal injection of d-luciferin potassium on day 1, 3, 5, 7, 9. **(d, e)** *In vivo* mRNA delivery efficacy of Pool 1-LNP 3 were measured by the IVIS imaging system and expression for 7 days, a two-tailed Student’s t-test data are represented as mean ± standard deviation, significance thresholds: \**P* < 0.05, ** *P* < 0.01, *** *P* < 0.001, **** *P* < 0.0001, n = 3 biologically independent mice per group.

### Delivery of *Msh5* mRNA using LNP3 restored spermatogenesis in *Msh5^D486Y/D486Y^* mice with spermatogenic disorders

To evaluate whether spermatogenic disorders could be rescued via the novel mRNA delivery strategy, *Msh5^D486Y/D486Y^* male mice with meiotic arrest as described as previously was chosen in the current study^8^. *Msh5* mRNA and *EGFP* mRNA were encapsuled in LNP3 with 1:1 mass ratio and injected to *Msh5^D486Y/D486Y^* male mice (Supplementary Fig 3a). Testicular wholemount fluorescence imaging on day 7, 14, and 21 after injection revealed the expression of EGFP sustained in the seminiferous tubules for more than 14 days (Supplementary Fig 3b). Hematoxylin and eosin staining revealed that while spermatogenesis was arrested at the spermatocyte stage in untreated *Msh5^D486Y/D486Y^* testis, elongated spermatids occurred in the seminiferous tubules by day 21 and disappeared on day 28 after delivery (Fig 3a). Furthermore, we performed immunofluorescence staining for germ cell markers of each stage, including SYCP3 (spermatocyte marker), AKAP3 (spermatid marker), TNP1 (elongating spermatid marker), and peanut agglutinin (PNA)-lectin to label the acrosome of the spermatid. The results confirmed the presence of PNA-positive round spermatids as early as day 7 post-treatment, albeit in low numbers (Supplementary Fig 3c). By day 14 AKAP3-positive spermatids had increased significantly in the seminiferous tubules and by day 21 most spermatids in the tubules had completed spermiogenesis (Fig 3b, c and Supplementary Fig 3d). However, elongated spermatids were absent in the testes collected on days 28 and 35 after injection, indicating transient expression of the mRNA within spermatocytes, sufficient to facilitate meiotic progression (Fig 3b, c).

**Fig 3.**
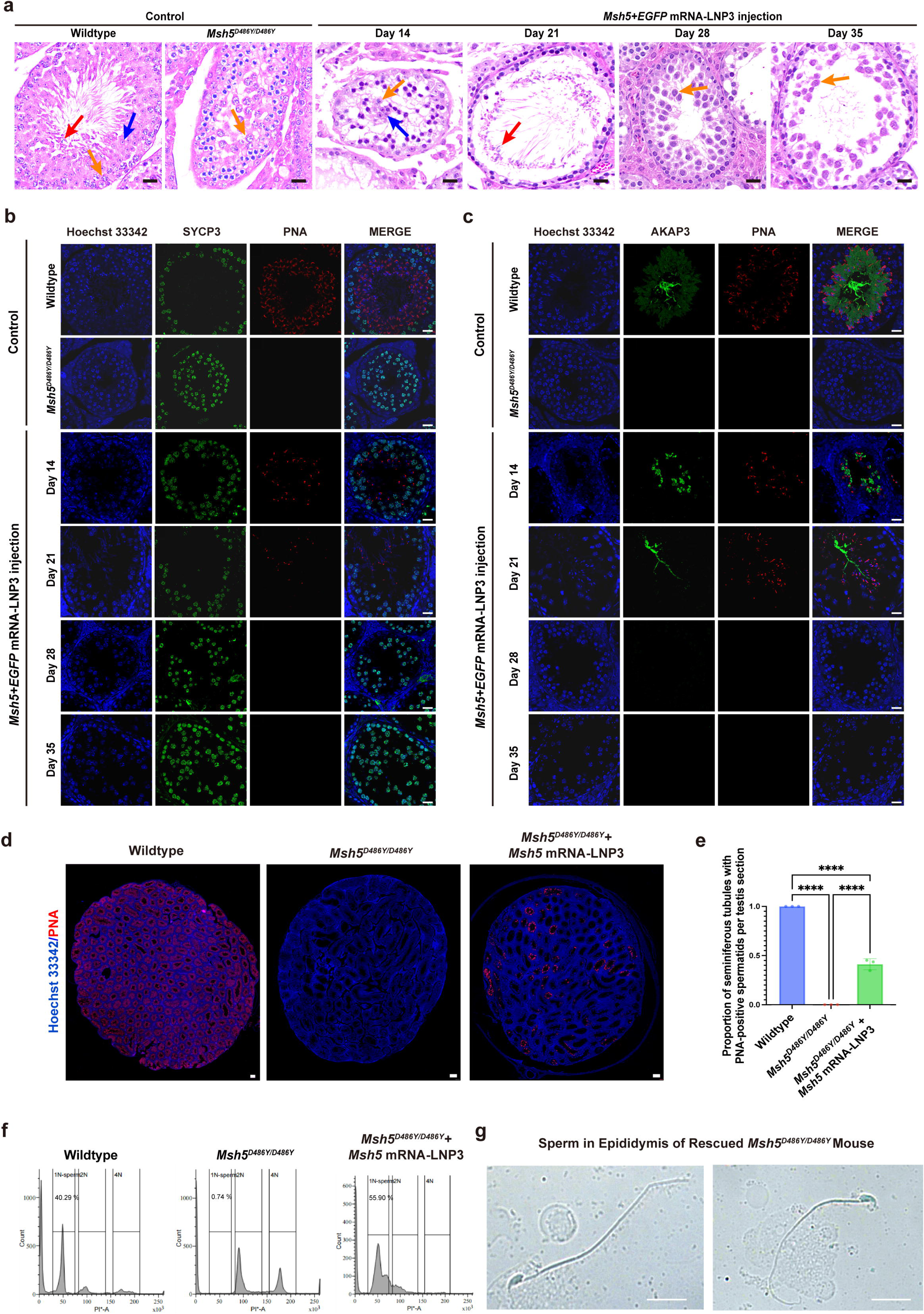
The restoration of spermatogenesis in *Msh5^D486Y/D486Y^* male mice. **(a)** H&E staining of *Msh5^D486Y/D486Y^* testis sections harvested on day 14, 21, 28, and 35 after injection of *Msh5* mRNA+*EGFP* mRNA-LNP3 revealed the emergence of elongated spermatids in day 21 and diminished in day 28, yellow arrow indicated spermatocytes, blue arrow indicated round spermatids, red arrow indicated elongated spermatids, scale bar = 20 μm. **(b, c)** Immunofluorescence staining showed the expression of SYCP3, PNA, and AKAP3 in *Msh5^D486Y/D486Y^*testis sections harvested on day 14, 21, 28, and 35 after injection of *Msh5* mRNA+*EGFP* mRNA-LNP3, scale bar = 20 μm. **(d)** Immunofluorescence staining showed the expression of PNA in whole testicular section from wildtype, *Msh5^D486Y/D486Y^* and *Msh5^D486Y/D486Y^*mice on day 21 after injection of *Msh5* mRNA+*EGFP* mRNA-LNP3, scale bar = 100 μm. **(e)** Comparison of the proportion of PNA positive seminiferous tubules in wildtype, *Msh5^D486Y/D486Y^*and *Msh5^D486Y/D486Y^* rescued testis, data are represented as mean ± standard deviation, n = 3 biologically independent mice per group, one-way ANOVA and Tukey’s multiple comparisons test. \*\*\*\**P* < 0.0001. **(f)** Comparison of the proportion of haploid germ cells by flow cytometry staining with propidium iodide (PI). **(g)** Morphology of sperm from epididymis in day 28 after injection, scale bar = 10 μm.

To evaluate the therapeutic rescue efficiency of mRNA-LNP3 delivery, we quantified seminiferous tubule with PNA-positive spermatids at 21 days after injection. The results revealed that 31.91±2.97% of tubules cross-sections exhibited PNA-positive spermatids within their luminal compartments, indicating restored spermatogenesis in these seminiferous tubules (Fig 3d, e). Flow cytometric quantification of testicular cell suspensions showed 55.9% of the testicular cells were haploid germ cells after mRNA delivery (1N population), demonstrating significant rescue efficiency compared with untreated *Msh5^D486Y/D486Y^* testis (Fig 3f). Furthermore, cytologic examinations revealed that spermatozoa in the epididymis showed normal head condensation, intact acrosomal structures, and typical flagellar architecture (Fig 3g).

In order to delineate the critical window for presence of elongated spermatids, testis and epididymides were collected on days 14, 16, 18, 20, 22, 24, 26 and 28 after mRNA delivery. H&E and immunofluorescence staining analysis indicated the presence of elongated spermatids in the seminiferous tubules by day 18 and were no longer detectable by day 22 (Fig 4a, c, d and Supplementary Fig 5), whereas H&E staining analysis revealed that there were no spermatozoa until day 28 after injection in the epididymis (Fig 4b). The percentages of PNA-positive seminiferous tubules at 14, 16 and 20 days after injection were 32.89±1.44%, 32.91±3.57% and 33.47±2.94% respectively (Fig 4e-g).

**Fig 4.**
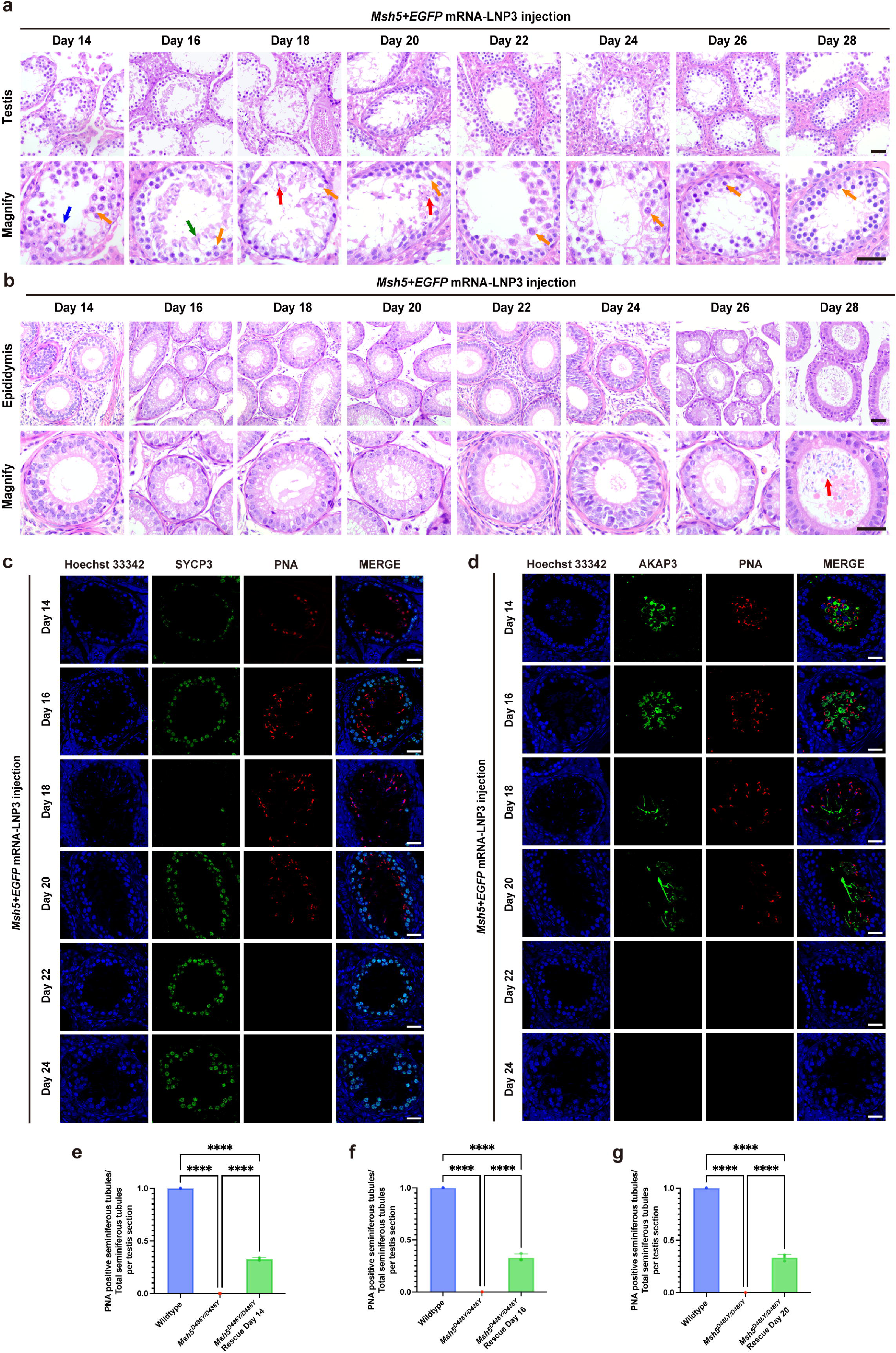
Timeline of restoration of spermatogenesis in *Msh5^D486Y/D486Y^* mice. **(a, b)** Hematoxylin and eosin staining of *Msh5^D486Y/D486Y^* testis and caput epididymis sections harvested in 14, 16, 18, 20, 22, 24, 26 and 28 days after injection of *Msh5* mRNA+*EGFP* mRNA-LNP3, yellow arrow indicated spermatocytes, blue arrow indicated round spermatids, green arrow indicated elongating spermatids and red arrow indicated elongated spermatids, scale bar = 50 μm. **(c, d)** Immunofluorescence staining showed the expression of SYCP3, AKAP3, and PNA in *Msh5^D486Y/D486Y^*testis sections harvested in 14, 16, 18, 20, 22 and 24 days after injection of *Msh5* mRNA+*EGFP* mRNA-LNP3, scale bar = 20 μm. **(e, f, g)** Comparison of the proportion of PNA positive seminiferous tubules in wildtype, *Msh5^D486Y/D486Y^*and *Msh5^D486Y/D486Y^* rescued testis post injection day 14, 16 and 20, data are represented as mean ± standard deviation, n = 3 biologically independent mice per group, one-way ANOVA and Tukey’s multiple comparisons test. \*\*\*\**P* < 0.0001.

To evaluate whether the delivered mRNA integrated into rescued sperm genome, we collected the sperms from testis on day 20 after injection and performed genotyping on 10 randomly selected sperms (Fig 5a), Sanger sequencing revealed that all 10 sperm samples exhibited mutation genotype (*Msh5* p.D486Y) (Fig 5b, c). To further evaluate the fertility of the mice after *Msh5* mRNA-LNP3 delivery, we microinjected 53 sperms into wildtype female mouse oocytes and observed 48 embryos developed to the 2□cell stage on day 2 after intracytoplasmic sperm injection (ICSI). On day 5-6 after ICSI, we observed 37 embryos proceeded to morula stage and 30 embryos proceeded to blastocysts (Fig 5d). Sanger sequencing revealed that all 10 randomly chosen blastocysts exhibited heterozygous genotypes (*Msh5* p.D486Y) (Fig 5e, f). After transplantation of the embryos, a live offspring was successfully delivered (Fig 5g). This genetic pattern provides conclusive evidence that the therapeutic mRNA delivery restored the full development of spermatogenesis in *Msh5^D486Y/D486Y^* male mice without genomic editing, highlighting the therapeutic potential of mRNA-based approaches in the treatment of spermatogenic disorders with genetic defects.

**Fig 5.**
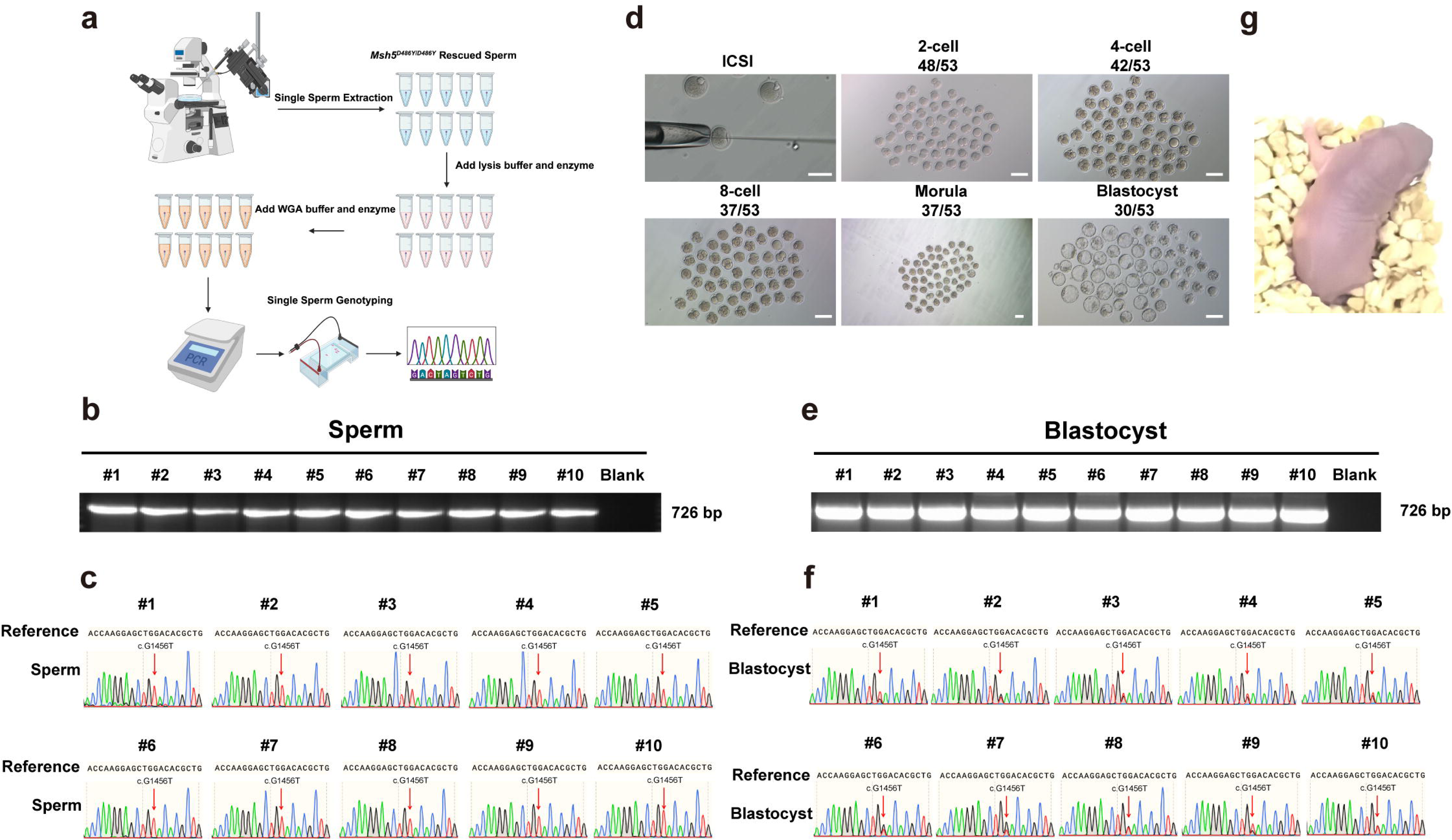
*Msh5* mRNA-LNP3 restored fertility in *Msh5^D486Y/D486Y^* mice and produced offspring. **(a)** Schematic diagram of single sperm genotyping of *Msh5* mRNA-LNP3 rescued sperm, (Created by Biorender.com). **(b, c)** Genotyping of the randomly chosen 10 single sperm from rescued testis, Sanger sequencing confirmed the variant of *Msh5* (c.G1456T). **(d)** Representative images of two-cell embryos, four-cell embryos, eight-cell embryos, morula embryos and blastocyst embryos after ICSI of the *Msh5* mRNA-LNP3 rescued sperm, scale bar =100 μm. **(e, f)** Genotyping of the randomly chosen 10 blastocysts, Sanger sequencing confirmed the heterozygous variant of *Msh5* (c.G1456T) in the blastocysts. **(g)** Offspring (F1) obtained using sperm from rescued *Msh5^D486Y/D486Y^* male mice by ICSI.

### Delivery of *Msh5* mRNA using LNP3 rescued the meiotic defects in *Msh5****^D486Y/D486Y^*** mice

To decipher the mechanism of restoration of spermatogenesis in *Msh5^D486Y/D486Y^*mice after *Msh5* mRNA LNP3 delivery, meiotic chromosomal spread analysis was performed to evaluate the meiotic progression and recombination. In wildtype DMC1 foci were abundant during early prophase I and progressively diminished in pachytene, indicating timely strand invasion and homologous recombination progression. In contrast, DMC1 foci of *Msh5^D486Y/D486Y^* spermatocytes persisted into pachytene, suggesting defects in strand invasion intermediate resolution and a prolonged recombination process. However, after injection of mRNA-LNP3, the rescued spermatocytes showed normal DSB formation and repair process by timely diminished DMC1 foci in pachytene (Supplementary Fig 4a). Furthermore, the result of chromosomal spread analysis by SYCP1 and SYCP3 indicated both of the *Msh5^D486Y/D486Y^*and the rescued spermatocytes could form normal synaptonemal complex (Supplementary Fig 4b).

The mismatch repair protein MLH1 was used to mark the formation of crossovers during meiotic recombination^35^. In mid-pachytene spermatocytes, MLH1 foci appeared as ∼22 discrete foci per nucleus, with at least one per homolog pair, evenly distributed along the synapsed axes^36^. The result of chromosomal spread analysis showed almost no MLH1 foci in *Msh5^D486Y/D486Y^* pachytene spermatocytes which is consistent with previous studies^8^, indicating the elimination of crossovers. In contrast, the rescued group displayed a significantly increased number of MLH1 foci, suggesting successful restoration of crossover formation (Supplementary Fig 4c). Altogether, these results suggested that delivery of *Msh5* mRNA LNP3 could rescue the DSB combination defects and meiotic arrest in *Msh5^D486Y/D486Y^* pachytene spermatocytes.

### Safety of *Msh5* mRNA-LNP3 delivery via rete testis microinjection *in vivo*

To confirm the safety of *Msh5* mRNA-LNP delivery *in vivo*, we tested the safety of Pool1-LNP3 and *Msh5* mRNA *in vivo* respectively. Firstly, we delivered PBS, 2 μg and 4 μg *EGFP* mRNA-LNP3 into seminiferous tubules of wildtype mice via rete testis microinjection to explore the safety of LNP. And multiple tissues throughout the body were collected, including the testis, blood, heart, liver, spleen, lungs, kidneys and brain after 24 hours. Gross photograph showed no significant abnormalities in any organs assessed (Fig 6a). Also, immunofluorescence analysis revealed that *EGFP* mRNA did not leak into other major organs throughout the body after rete testis injection (Fig 6b). Western blot revealed that EGFP was expressed in testis after mRNA-LNP delivery via concentration-dependent manner, and no expressions of EGFP except testis were detected in multiple tissues throughout the body (Fig 6c, d). Furthermore, H&E staining revealed no significant abnormalities in any organs assessed (Fig 6e). Elisa assay showed there were no significant differences of the expression of pro-inflammatory cytokines (IL-1β) among the three group (PBS, 2 μg and 4 μg, respectively) (Fig 6f).

**Fig 6.**
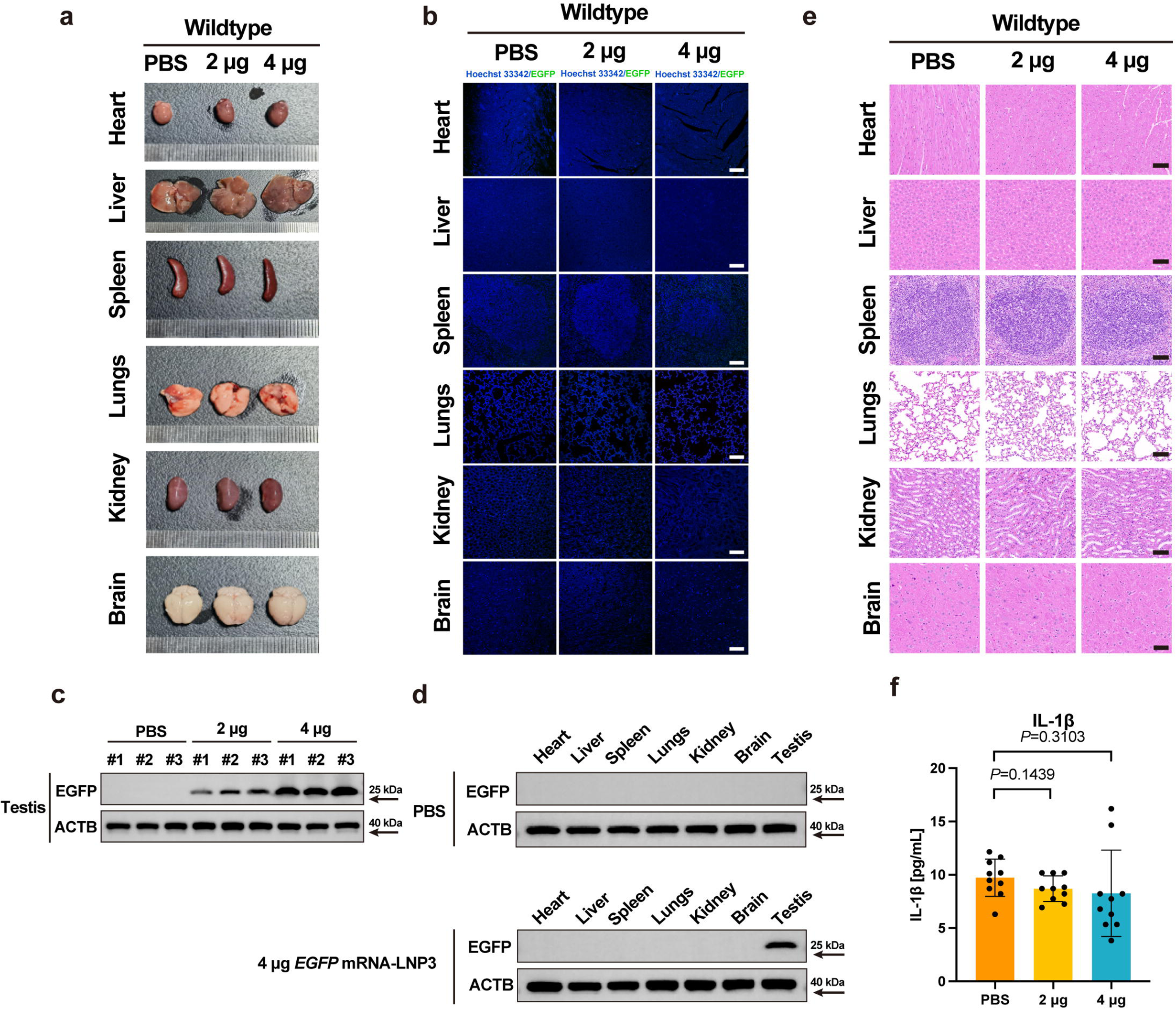
Systemic safety validation of LNP3 and payload. **(a)** No macroscopic lesions in major organs (heart, liver, spleen, lungs, kidney and brain tissue) following injection of varying *EGFP* mRNA-LNP3 doses (2 μg/per testis and 4 μg/per testis) in wildtype mice, samples were collected 24 h after injection, n = 3 biologically independent mice per group. **(b)** Immunofluorescence staining revealed no ectopic expression of EGFP in major organs (heart, liver, spleen, lungs, kidney and brain tissue) following injection of varying *EGFP* mRNA-LNP3 doses (2 μg/per testis and 4 μg/per testis) in wildtype mice, scale bar = 50 μm, n = 3 biologically independent mice per group. **(c)** Western blot analysis showed the expression of EGFP in testicular samples from wildtype testis following injection of varying *EGFP* mRNA-LNP3 doses (2 μg/per testis and 4 μg/per testis) and PBS as negative control, n = 3 biologically independent mice per group. **(d)** Western blot analysis showed the expression of EGFP in major organ samples (heart, liver, spleen, lungs, kidney and brain tissue) from wildtype mice following injection of *EGFP* mRNA-LNP3 high doses 4 μg/per testis and PBS, n = 3 biologically independent mice per group. **(e)** Hematoxylin and eosin staining analysis showed preserved tissue architecture without significantly pathological alterations in major organs (heart, liver, spleen, lungs, kidney, and brain), scale bar = 50 μm, n = 3 biologically independent mice per group. **(f)** Elisa assay showed the expression of IL-1β in the blood after injection of varying *EGFP* mRNA-LNP3 doses (2 μg/per testis and 4 μg/per testis), data are represented as mean ± standard deviation, n = 10 biologically independent mice per group, one-way ANOVA and Tukey’s multiple comparisons test.

Subsequently, we evaluated the safety of *Msh5* mRNA *in vivo* via rete testis microinjection of mixture of *Msh5* mRNA and *EGFP* mRNA at 2 μg in *Msh5^D486Y/D486Y^* male mice. Multiple tissues throughout the body on day 21 after injection were collected, including heart, liver, spleen, lungs, kidney and brain. And the histological analysis showed that there were no significant abnormalities in any organs assessed compared with untreated *Msh5^D486Y/D486Y^* male mice (Supplementary Fig 6). Collectively, these results illustrated that *Msh5* mRNA-LNP3 delivery via rete testis microinjection *in vivo* showed no obvious toxicity in multiple tissues throughout the body.

### mRNA-LNP delivery restored spermatogenic disorders in *Maps* KO Mice: toward versatile therapy

To explore the broad applicability of mRNA-LNP technology, *Maps* (*C3orf62*) KO mice via CRISPR/Cas9 mediated genome editing was generated as described previously^30^. *Maps* KO mice faithfully recapitulated the human infertility phenotype as we reported previously, showing complete spermatogenic arrest at the spermatocyte stage. Intriguingly, sequential restoration of spermatogenesis was observed after *Maps* mRNA+*EGFP* mRNA LNP3 (2 μg per testis) delivery via rete testis microinjection. Specifically, seminiferous tubules exhibited occurrence of round spermatids on day 14 after injection and elongated spermatids on day 21 after injection (Fig 7a). The results of immunofluorescence staining also revealed the PNA-positive round spermatids and AKAP3-positive elongating spermatids could be detected on day 14 post injection and TP1 positive elongated spermatids emerged on day 21 (Fig 7b, c and Supplementary Fig 7). The percentage of PNA-positive seminiferous tubules at 21 days after injection was about 29.47±3.2% (Fig 7d). Also, a comprehensive safety evaluation was conducted in *Maps* KO mice after mRNA delivery through histopathological analysis of multiple tissues throughout the body. Tissue specimens including heart, liver, spleen, lungs, kidney, and brain were systematically collected from both untreated *Maps KO* controls and rescued mice at 21 days after injection. Rigorous histological examination revealed no significant pathological alterations observed in the rescued tissues compared with *Maps* KO mice (Fig 7e). These findings demonstrated no obvious systemic toxicity associated with mRNA delivery in these mice models. The successful rescue of meiotic arrest in these models validate the platform’s potential for broad applicability in NOA-affected patients with genetic disorders.

**Fig 7.**
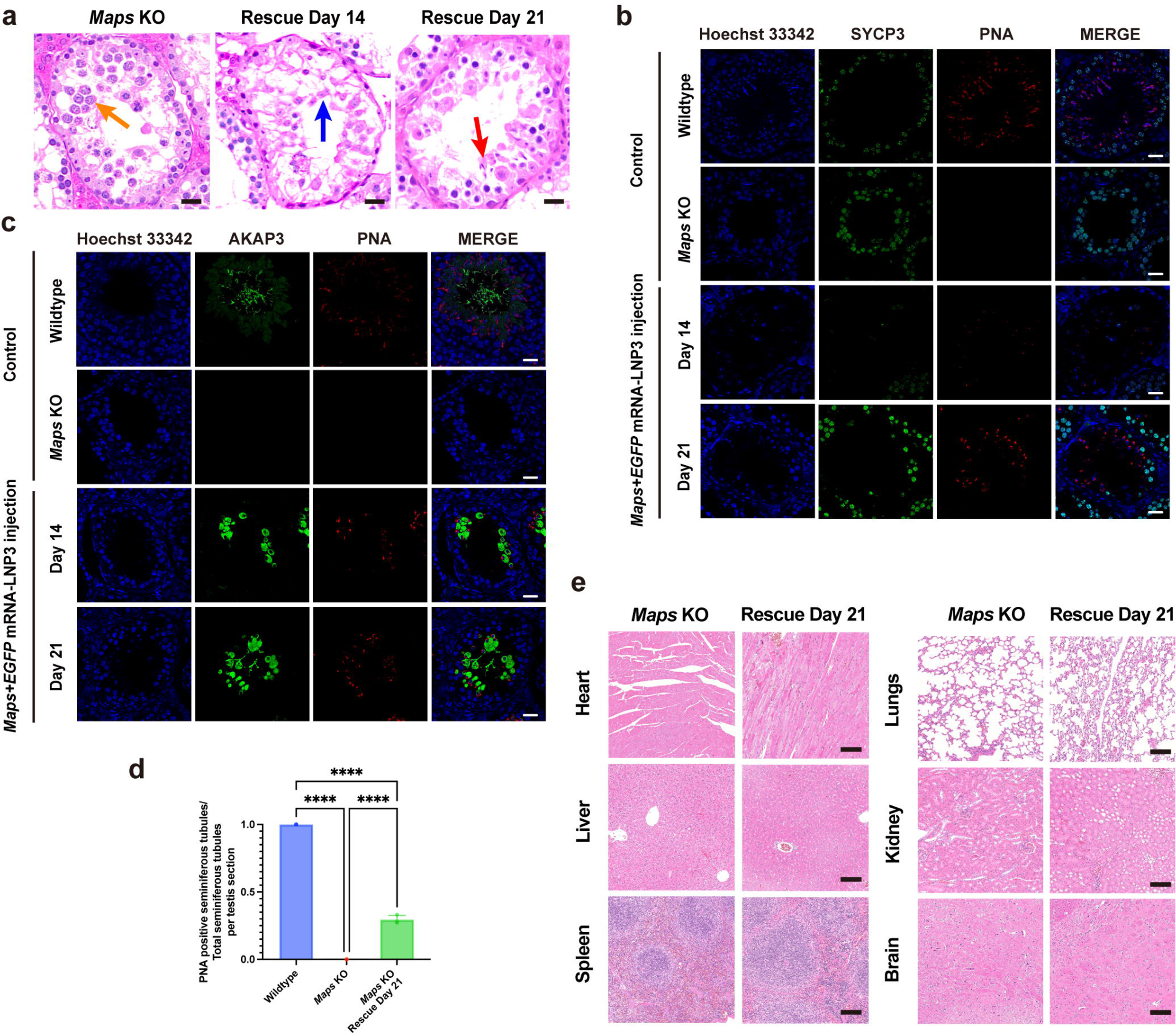
Restoration of spermatogenesis in *Maps* KO male mice. **(a)** Hematoxylin and eosin staining revealed the appearance of round spermatids and elongated spermatids on day 14 and 21 after injection respectively, yellow arrow indicated spermatocytes, blue arrow indicated round spermatids, red arrow indicated elongated sperm, scale bar = 20 μm. **(b, c)** Immunofluorescence staining showed the expression of SYCP3, PNA, and AKAP3 in *Maps* KO testis sections harvested on day 14 and 21 after injection of *Maps* mRNA+*EGFP* mRNA-LNP3, scale bar = 20 μm. **(d)** Comparison of the proportion of PNA positive seminiferous tubules in wildtype, *Maps* KO and *Maps* KO rescued testis at day 21after injection, data are represented as mean ± standard deviation, n = 3 biologically independent mice per group, one-way ANOVA and Tukey’s multiple comparisons test. \*\*\*\**P* < 0.0001. **(e)** Hematoxylin and eosin staining analysis showed preserved tissue architecture without significantly pathological alterations in major organs (heart, liver, spleen, lungs, kidney, and brain) in *Maps* KO mice at 21 days after injection, scale bar = 50 μm, n = 3 biologically independent mice per group.

## Discussion

In the current study, we developed a novel therapy for genetic spermatogenic disorders. Herein, a spermatocytes-tropic LNP (Pool1-LNP3) was identified via a library of 30 ionizable lipids screening. And it was revealed that *in vivo* delivery of mRNA LNP3 could restore spermatogenesis in *Msh5^D486Y/D486Y^* and *Maps* KO mouse models with meiotic arrest. Notably, the offspring without genomic integration was born using intracytoplasmic sperm injection (ICSI) derived from rescue of *Msh5^D486Y/D486Y^* mouse and embryo transfer. Furthermore, no obvious inflammation and histologic damage in multiple tissues throughout the body were detected after *in vivo* delivery of LNP. Thus, we developed a novel testicular mRNA-LNP delivery system as an effective solution for genetic spermatogenic disorders.

Prior attempts to address spermatogenic disorders with genetic defects have faced substantial limitations. Current therapeutic strategies for restoration of spermatogenesis in mice models with spermatogenic disorders derived from clinical NOA with genetic variants involved two CRISPR-Cas9-based approaches, including *ex vivo* editing of spermatogonial stem cells (SSCs) followed by testicular transplantation^37–39^, and *in vivo* CRISPR-Cas9 delivery into seminiferous tubules for genetic correction^8^. Both strategies can restore spermatogenesis by correcting pathogenic variants, however, their clinical applications faced insurmountable barriers. Persistent concerns regarding CRISPR-Cas9 off-target effects and incomplete validation of long-term safety *in vivo* ^40–42^. Notably, genome editing of human gametes and zygotes was strictly prohibited, rendering these approaches clinically inapplicable. Compounding these challenges, the absence of robust protocols for long-term human SSCs culture *in vitro* further limits feasibility^43,44^. In the current study, the novel mRNA-LNP platform circumvents these limitations through a genomic non-integrative mechanism. By delivering functional mRNA directly to spermatocytes, they transiently translated to wild-type proteins in cytoplasm, thus this approach achieves therapeutic protein expression without altering genomic integrity—a critical safety advantage over permanent genome editing.

RNA is a negatively charged molecule, and naked mRNA is unable to cross the cell membrane directly due to its charge properties, unless facilitated by techniques such as electroporation. However, while electroporation can enhance RNA delivery into cells, it induces obvious cellular damage and poses challenges for clinical applications. Additionally, RNA is highly susceptible to degradation by nucleases, which compromises its stability *in vivo*. Therefore, appropriate carriers should be used in the mRNA-based therapeutic delivery due to its efficient delivery and biological stability. Previous studies have revealed that AAV vectors could be used to rescue spermatogenic disorders in mouse via delivery of therapeutic plasmids into seminiferous tubules^45,46^. However, the clinical applications of AAV-based therapies faced critical limitations in reproductive medicine. First, inherent risks of genomic integration, albeit at low frequencies, were identified in AAV-mediated plasmid delivery system, raising concerns about insertional mutagenesis in germ cells^47,48^. Secondly, pre-existing AAV DNA has been identified in human seminal plasma. It was detected in 35.9% (28/78) of men with abnormal semen parameters in a clinical cohort. This result indicated that AAV-mediated plasmid delivery may compromise therapeutic efficacy through immune clearance, also suggesting potential associations between viral exposure and impaired spermatogenesis^49^. At last, AAV’s constrained payload capacity (∼4.7 kb) limits its utility for delivery of large genes^45^. In contrast, our LNP platform addresses these challenges comprehensively. As a non-viral delivery system, LNPs eliminate risks of genomic integration while circumventing AAV-related immunogenicity. Also, the safety of the mRNA delivery system was validated by the global administration of billions of LNP-based COVID-19 mRNA vaccines^50,51^. The modular design of LNPs accommodates larger mRNA payloads, enabling delivery of full-length therapeutic transcripts without size constraints. Furthermore, LNPs bypassed the complex manufacturing requirements of viral vectors, significantly enhancing scalability for clinical applications.

Current advancements in protein replacement therapies encompass diverse RNA modalities, including conventional mRNA, circular RNA (circRNA), and self-amplifying RNA (saRNA). Notably, previous studies demonstrated that LNP with the cargo (saRNA encoding *DMC1*) could rescue spermatogenic arrest in *Dmc1* KO mice. The saRNA systems achieved prolonged intracellular protein expression through RNA replication (∼1 month). However, it was engineered with non-structural protein elements derived from alphaviruses, raising concerns about the safety. Furthermore, endogenous *Dmc1* expression in wild-type mice is tightly regulated, with mRNA and protein predominantly localized to spermatocytes during meiosis and absent in post-meiotic spermatids. This pattern is conserved across key meiotic regulators in mammals, suggesting advantage in safety of transient rather than sustained expression for rescue of spermatogenic disorders ^52^.

Additionally, rete testis microinjection was used to topical medication in our mRNA LNP delivery system, which facilitated the localization of mRNA-LNPs in the lumen of seminiferous tubule. Benefiting from the physiological characteristics of the blood-testis barrier (BTB), the probability of mRNA-LNPs spillover into systemic circulation is exceedingly low. This strategy is therefore designed to minimize ectopic expression in tissues beyond the testis. The rete testis microinjection method not only ensures the localized delivery of mRNA-LNPs into the seminiferous tubules but also avoids the testicular tissue damage. Consequently, this technique holds greater potential for future clinical applications.

Although our study has demonstrated promising therapeutic efficacy in murine models, several limitations must be addressed to advance clinical application. On the one hand, comprehensive validation of the platform’s broad-spectrum applicability requires testing in mouse models with different genetic defects. On the other hand, while our study identified the spermatocyte-tropic LNP which could address meiotic arrest with genetic defects, spermatogenic arrest at spermatogonial or spermatid stages with genetic variants may necessitate distinct LNP delivery strategies. Further studies should be performed to develop spermatogonial stem cell/spermatid-targeting platforms through either high-throughput ionizable lipid screening (passive targeting) or antibody-conjugated active targeting.

Collectively, our findings identified a novel testicular mRNA-LNP delivery system as an effective solution for spermatogenic arrest with genetic defects. This evidence suggested broad clinical application potential of LNP-based mRNA therapy as a safe and versatile solution for NOA-affected patients harboring diverse genetic etiologies.

## Author contribution

C.W.Z. carried out the experiments, data analysis, assisted with the experimental design, and wrote the manuscript. Nan Liang assisted with the design of the vector and the animal experiments., W.B.L., S.X. and P.L. carried out the experiments and data analysis. W.Z.N., and S.H. helped with immunofluorescence staining. Na Li provided technical help in flow cytometry analysis. N.J.O., E.L.Z., R.H.T., Y.H.H. and F.J.Z. provided technical help in rete testis microinjection. Y.F.S., H.W.B., J.P.Z. and X.J.B. helped with animal experiments. D.W.Q. helped with data analysis and statistics. F.R.B. assisted with ICSI and embryo transplantation. C.C.Y., Z.L. and H.C. conceived the project, supervised all experiments and wrote and revised the manuscript. All authors fulfill the criteria for authorship.

## Completing of interest

C.C.Y., C.W.Z. and Z.L. have applied for a patent based on this study. All other authors declare no potential conflicts of interest.

## Acknowledgements

The authors acknowledge the *Msh5^D486Y/D486Y^* mice gifted by Pro. Fei Gao of Chinese Academy of Sciences and appreciate the help from Pro. Peng Wang, Dr. Yang Ji, Dr. Chen Zeng, Xingwu Liu and Bao Xiao of Southern University of Science and Technology and the help from Pro. Jun Zhang of Nanjing Medical University. We are appreciated the great help of Pro. Chong Li and Yijia Yang in the guidance of ICSI and embryo transplantation. We are also grateful to the entire team at Carcell BioPharma for their help by manufacturing LNP formulations.

---

**Fig 8.**
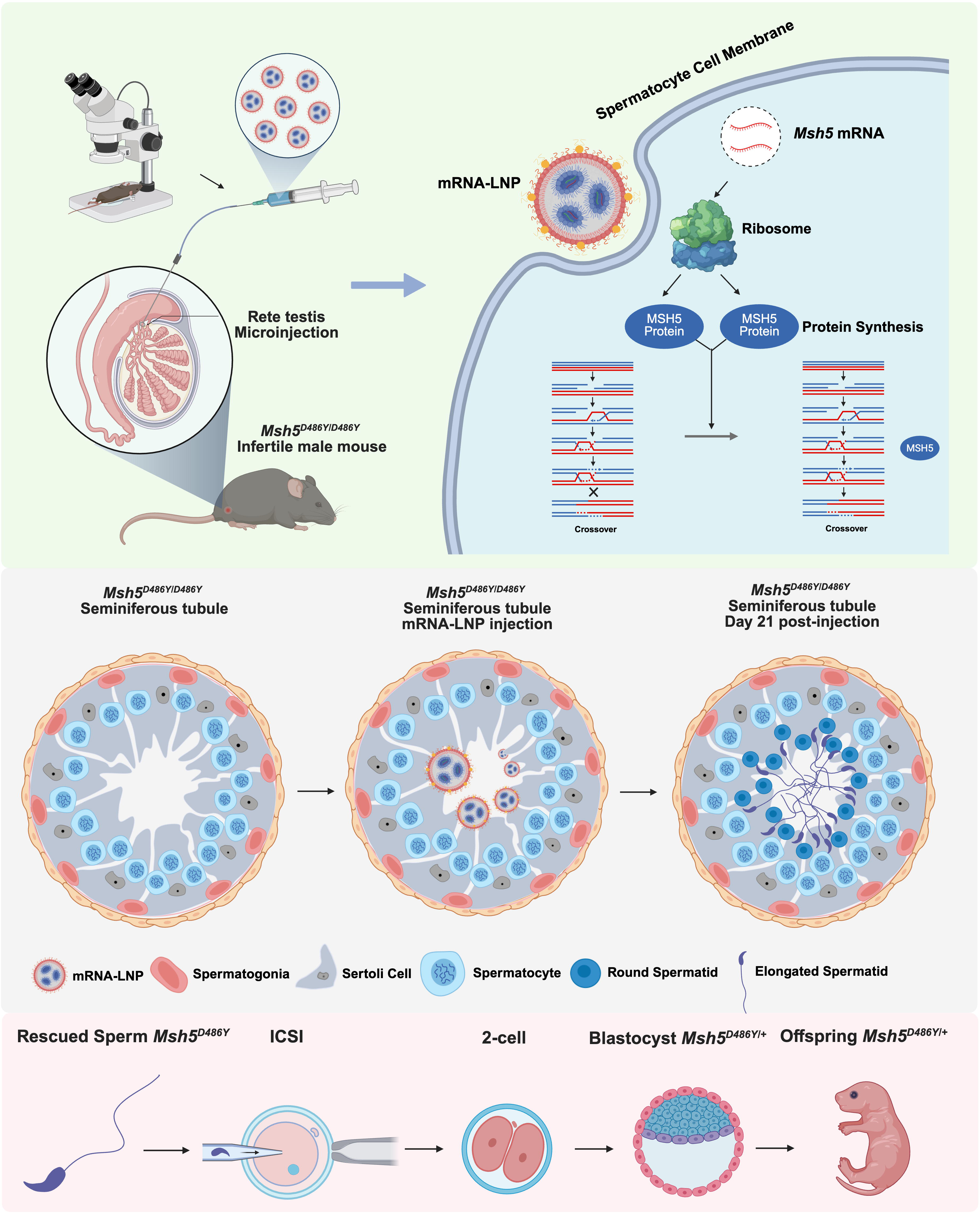
Schematic diagram of novel therapy for genetic spermatogenic disorder via testicular mRNA-LNP delivery.

**Fig S1.**
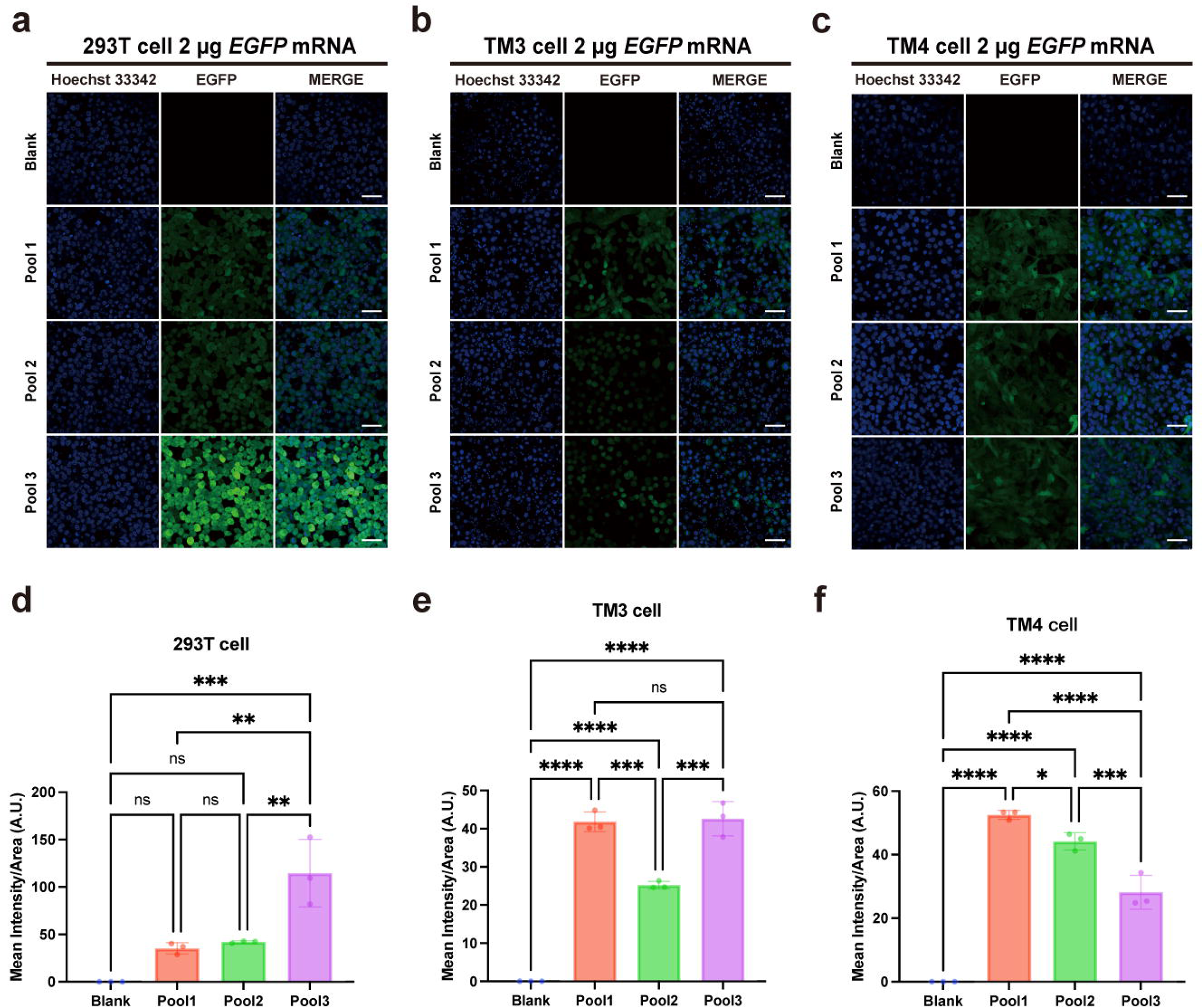
*In vitro* and *in vivo* validation of expression of *EGFP* mRNA-Pool 1, 2 and 3. **(a, b, c)** Expression of EGFP after 2 μg transfection with *EGFP* mRNA-LNP Pool 1, 2 and 3 in 293T, TM3 and TM4 cells cultured in 6-well plates, scales bar = 20 μm. **(d, e, f)** Mean intensity of EGFP expression after 2 μg transfection with *EGFP* mRNA-LNP Pool 1, 2 and 3 in 293T, TM3 and TM4 cells cultured in 6-well plates, data are represented as mean ± standard deviation, one-way ANOVA and Tukey’s multiple comparisons test. \**P* < 0.05, \*\**P* < 0.01, \*\*\**P* < 0.001, \*\*\*\**P* < 0.0001.

**Fig S2.**
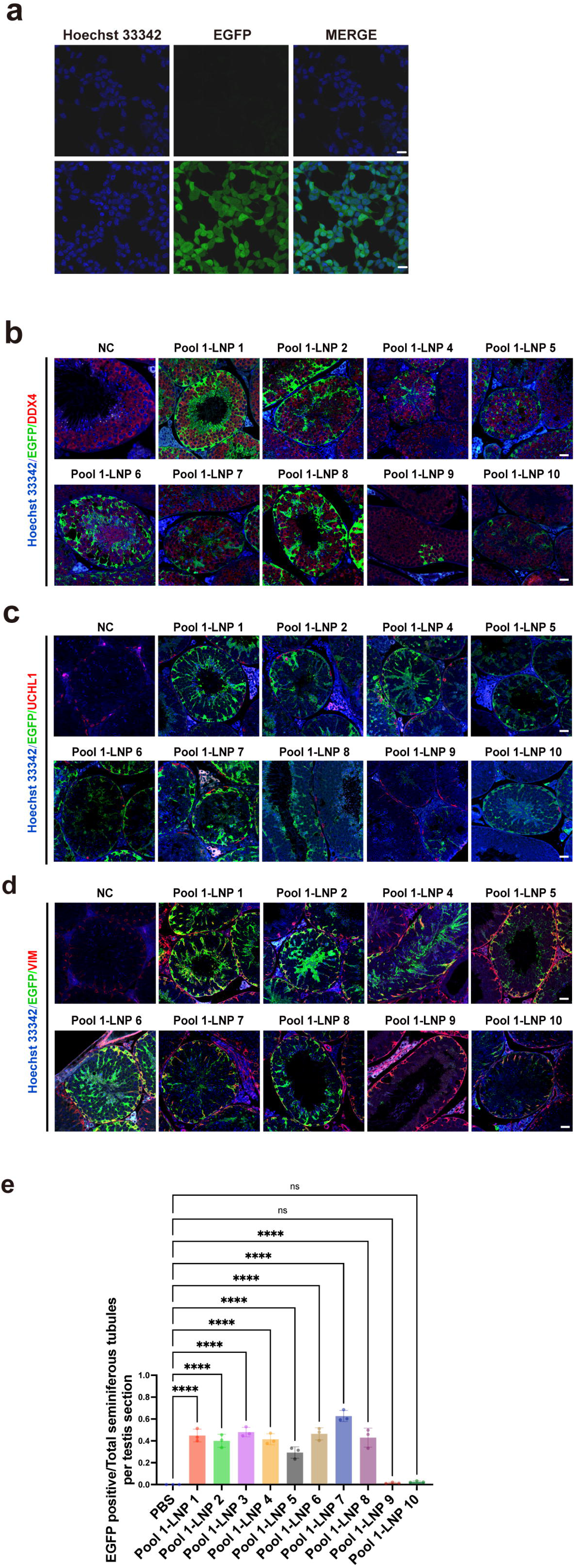
*In vivo* validation of expression of *EGFP* mRNA-LNP 1-10 in Pool1. **(a)** *In vitro* validation of expression of EGFP after 2 μg transfection with *EGFP* mRNA-Pool 1 LNP 3 in 293T cells cultured in 6-well plates. **(b, c, d)** Immunofluorescence staining showed the expression of EGFP and the germ cell marker DDX4, the spermatogonia marker UCHL1 and the Sertoli cell marker Vimentin in testicular sections of Pool 1 LNP 1-10 (without LNP 3) after injection, NC denoted the negative control group administered with PBS, scale bar = 20 μm. **(e)** The percentage of EGFP-positive seminiferous tubules relative to the total seminiferous tubule population, data are represented as mean ± standard deviation, n = 3 biologically independent mice per group, one-way ANOVA and Tukey’s multiple comparisons test. \*\*\*\**P* < 0.0001.

**Fig S3.**
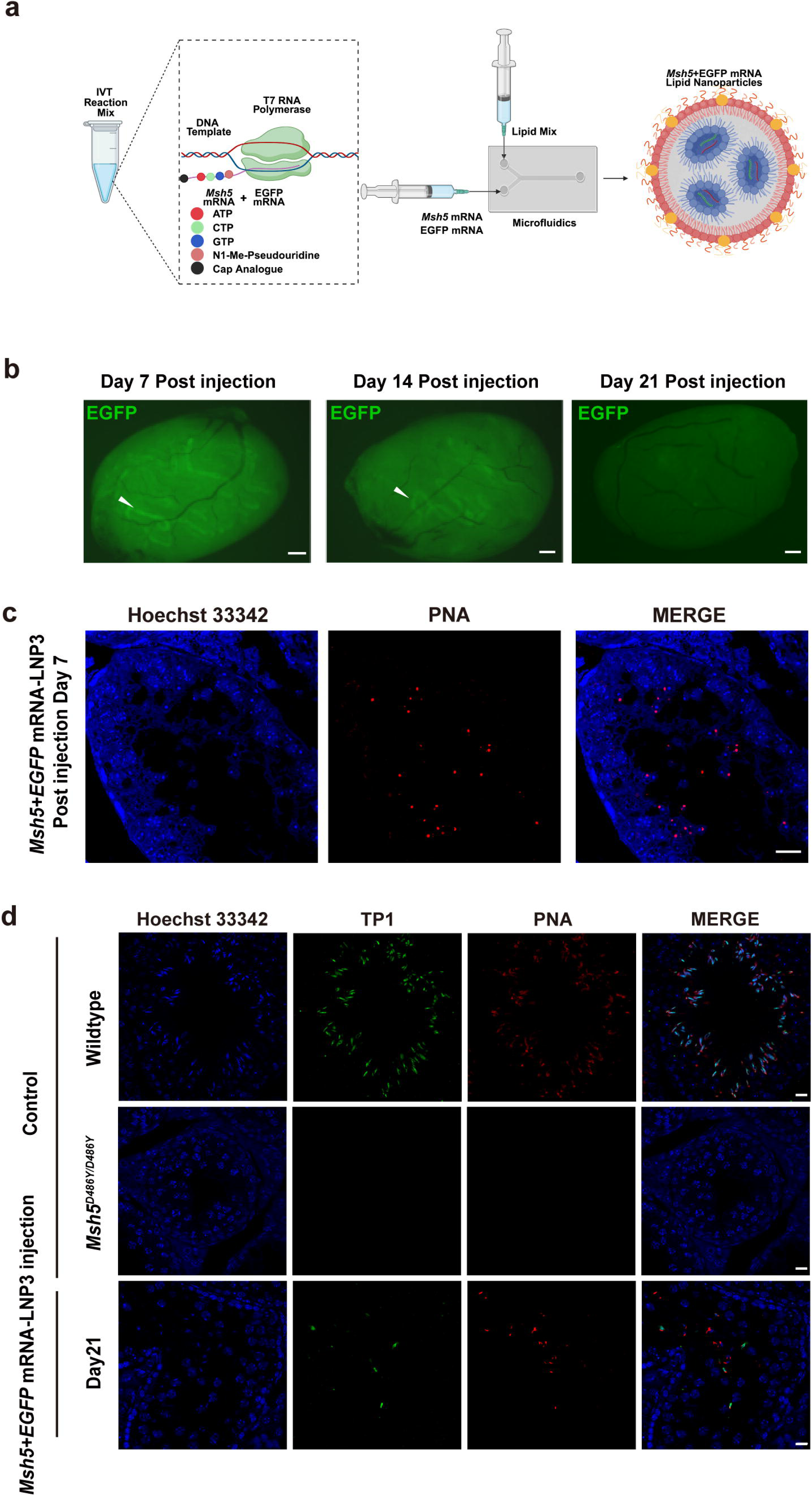
Recovery of spermatogenesis of *Msh5^D486Y/D486Y^*male mice. **(a)** Schematic diagram of *Msh5* mRNA + *EGFP* mRNA-LNP3 prepared by in vitro transcription and microfluidic systems. **(b)** Wholemount tissue fluorescence imaging of testis 7, 14 and 21 days after injection of *EGFP* mRNA-LNP3, white arrow head indicated the EGFP positive seminiferous tubules, scales bar = 5 mm. **(c)** Immunofluorescence staining showed PNA expression in *Msh5^D486Y/D486Y^* testis sections harvested in 7 days after injection of *Msh5* mRNA+*EGFP* mRNA-LNP3, scale bar = 20 μm. **(d)** Immunofluorescence staining showed the expression of PNA and TP1 in *Msh5^D486Y/D486Y^* testis sections harvested in 21 days after injection of *Msh5* mRNA+*EGFP* mRNA-LNP3, scale bar = 20 μm.

**Fig S4.**
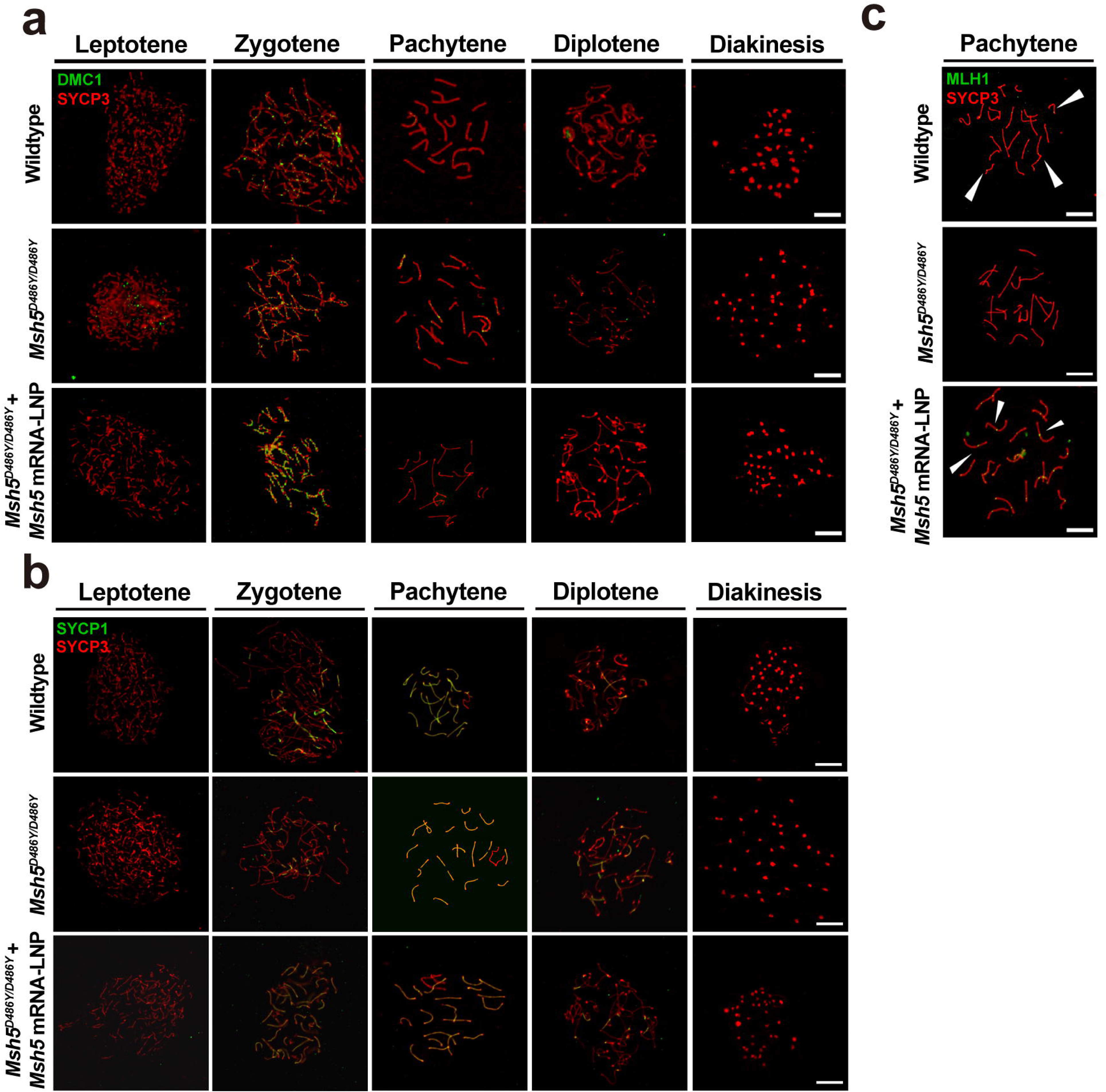
Chromosome spreading assay of *Msh5^D486Y/D486Y^* mice spermatocytes in day 3 post injection. **(a)** Chromosome spreads analysis showed the expression of DMC1 foci and SYCP3 in spermatocytes with different stages in wildtype mice, *Msh5^D486Y/D486Y^* mice and *Msh5* mRNA-LNP3 treated *Msh5^D486Y/D486Y^*mice. **(b)** Chromosome spreads analysis showed the expression of SYCP1 (green) and SYCP3 (red) spermatocytes from wildtype mice, *Msh5^D486Y/D486Y^* mice and *Msh5* mRNA-LNP3 treated *Msh5^D486Y/D486Y^* mice, scale bar = 10 μm. **(c)** Chromosome spreads analysis showed the expression of MLH1 foci in spermatocyte at pachytene stage of wildtype mice, *Msh5^D486Y/D486Y^* mice and *Msh5* mRNA-LNP3 treated *Msh5^D486Y/D486Y^* mice, scale bar = 10 μm.

**Fig S5.**
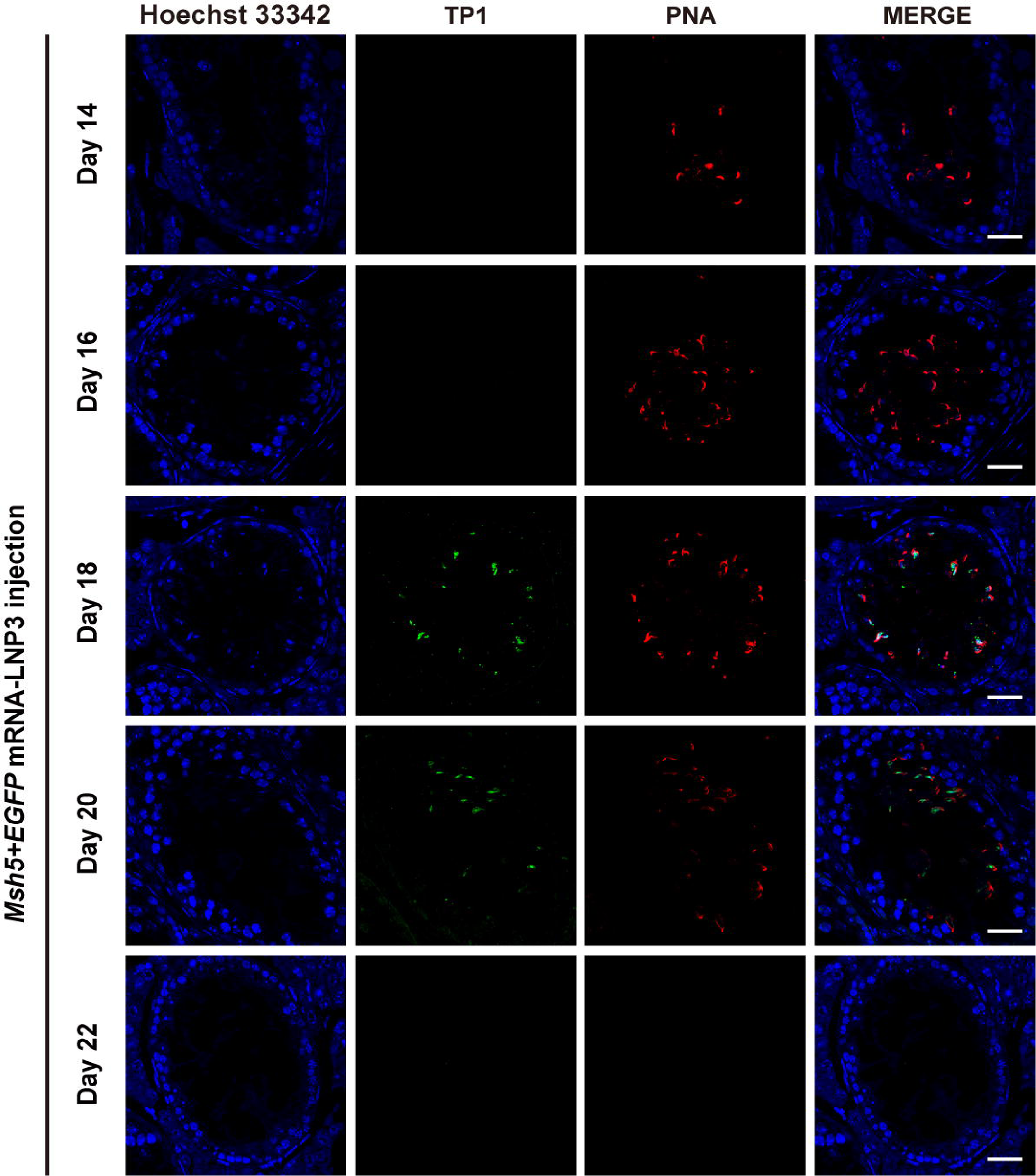
Timeline of rescued spermatids in *Msh5^D486Y/D486Y^* male mice after injection of *Msh5* mRNA+*EGFP* mRNA-LNP3. Immunofluorescence staining showed the expression of PNA and TP1 in *Msh5^D486Y/D486Y^* testis sections harvested in 14, 16, 18, 20 and 22 days after injection of *Msh5* mRNA+*EGFP* mRNA-LNP3, scale bar = 20 μm.

**Fig S6.**
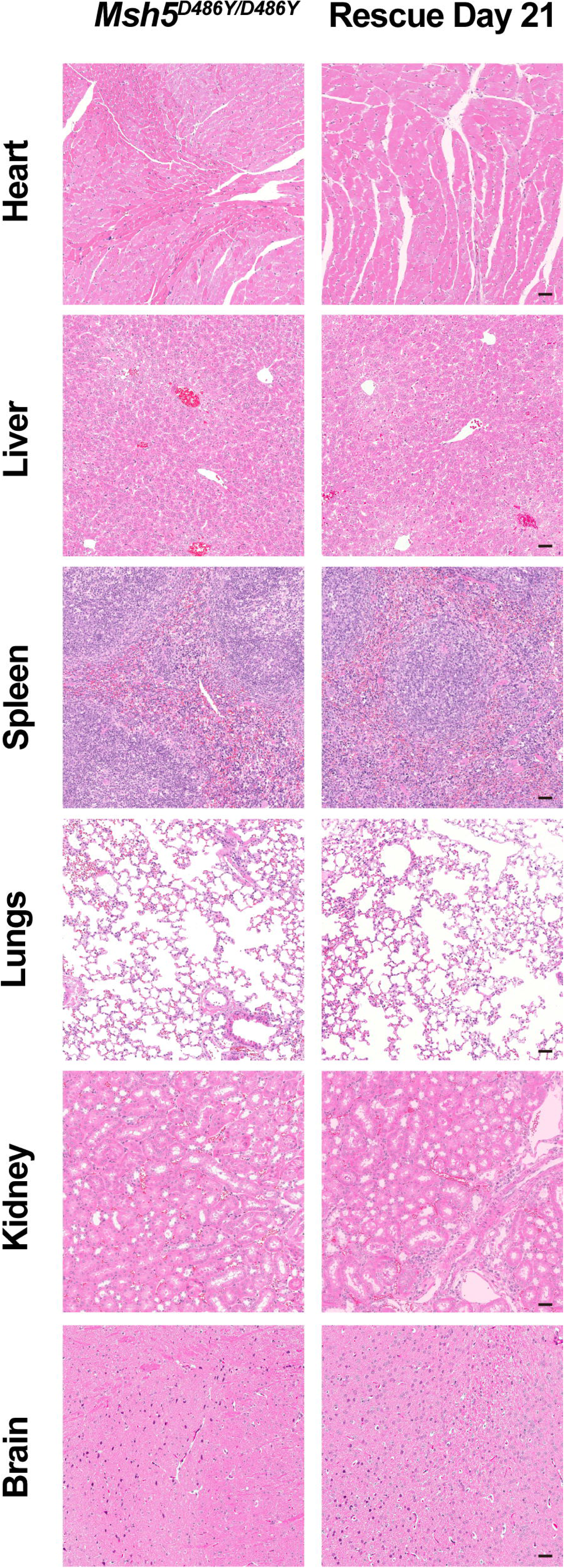
Safety validation of *Msh5* mRNA in Msh5^D486Y/D486Y^. Hematoxylin and eosin staining analysis showed revealed preserved tissue architecture without significantly pathological alterations in major organs (heart, liver, spleen, lungs, kidney, and brain) of *Msh5^D486Y/D486Y^* mice on day 21 after injection, scale bar = 50 μm.

**Fig S7.**
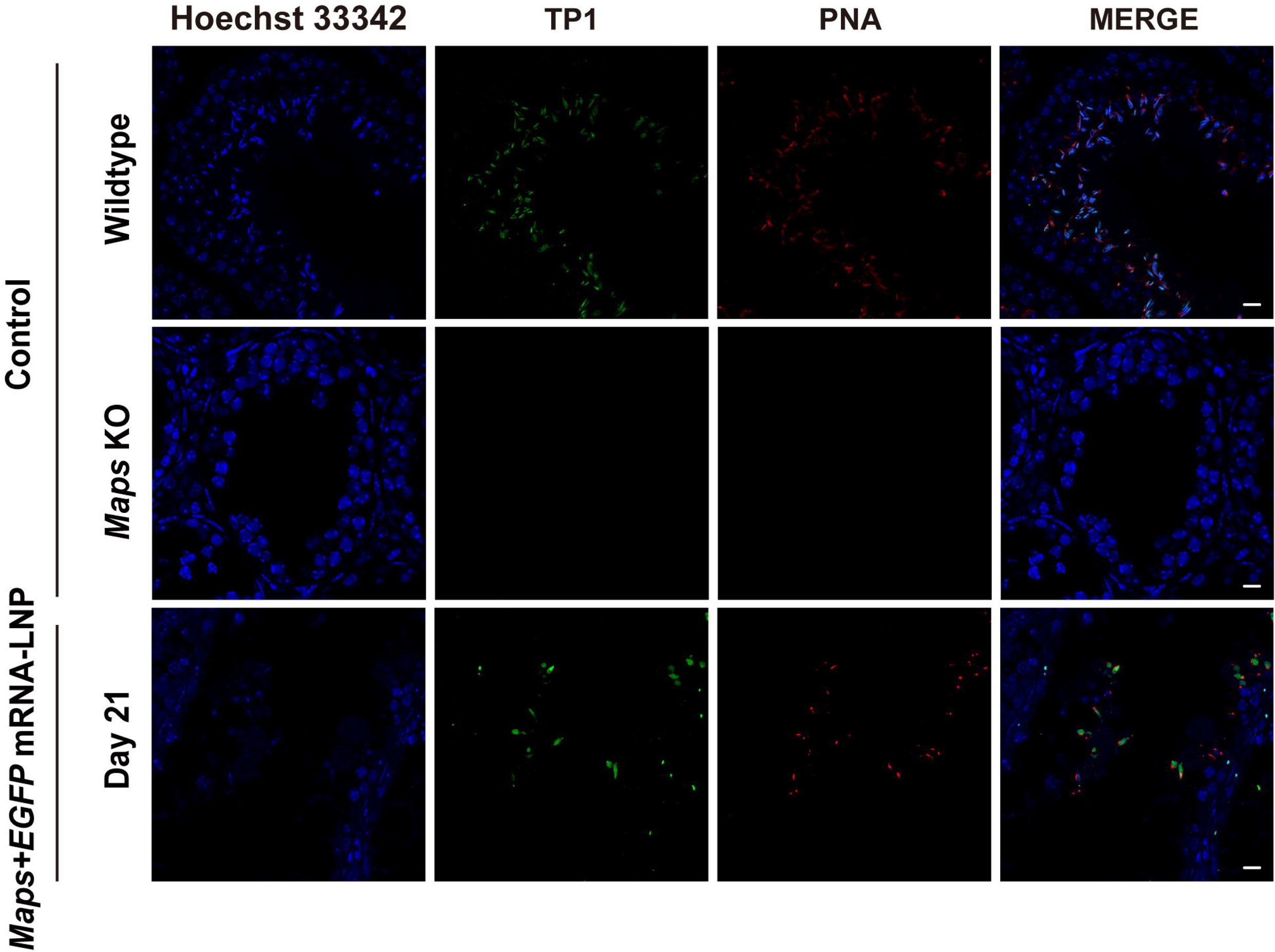
Recovery of spermatogenesis of *Maps* KO male mice. Immunofluorescence staining showed the expression of TP1 and PNA in *Maps* KO testis sections harvested in 21 days after injection of *Maps* mRNA+*EGFP* mRNA-LNP3, scale bar = 20 μm.

## Reference

1 Eisenberg, M. L., et al. Male infertility. Nat Rev Dis Primers 9, 49 (2023). 10.1038/s41572-023-00459-w

2 Agarwal, A., et al. Male infertility. Lancet 397, 319–333 (2021). 10.1016/S0140-6736(20)32667-2

3 Soumillon, M. et al. Cellular source and mechanisms of high transcriptome complexity in the mammalian testis. Cell Rep 3, 2179–2190 (2013). 10.1016/j.celrep.2013.05.031

4 Krausz, C. & Riera-Escamilla, A. Genetics of male infertility. Nat Rev Urol 15, 369–384 (2018). 10.1038/s41585-018-0003-3

5 Zhao, J. et al. Identification of a missense variant of MND1 in meiotic arrest and non-obstructive azoospermia. J Hum Genet 68, 729–735 (2023). 10.1038/s10038-023-01172-y

6 Zhang, Y. et al. Bi-allelic MEI1 variants cause meiosis arrest and non-obstructive azoospermia. J Hum Genet 68, 383–392 (2023). 10.1038/s10038-023-01119-3

7 Xu, J. et al. A homozygous frameshift variant in SYCP2 caused meiotic arrest and non-obstructive azoospermia. Clin Genet 104, 577–581 (2023). 10.1111/cge.14392

8 Chen, M. et al. Mutations of MSH5 in nonobstructive azoospermia (NOA) and rescued via in vivo gene editing. Signal Transduct Target Ther 7, 1 (2022). 10.1038/s41392-021-00710-4

9 Lin, Z. et al. The male pachynema-specific protein MAPS drives phase separation in vitro and regulates sex body formation and chromatin behaviors in vivo. Cell Rep 43, 113651 (2024). 10.1016/j.celrep.2023.113651

10 Kherraf, Z. E. et al. Whole-exome sequencing improves the diagnosis and care of men with non-obstructive azoospermia. Am J Hum Genet 109, 508–517 (2022). 10.1016/j.ajhg.2022.01.011

11 Sang, Q., Ray, P. F. & Wang, L. Understanding the genetics of human infertility. Science 380, 158–163 (2023). 10.1126/science.adf7760

12 Yang, C., Li, P. & Li, Z. Clinical application of aromatase inhibitors to treat male infertility. Hum Reprod Update 28, 30–50 (2021). 10.1093/humupd/dmab036

13 Bernie, A. M. et al. Outcomes of microdissection testicular sperm extraction in men with nonobstructive azoospermia due to maturation arrest. Fertil Steril 104, 569–573 e561 (2015). 10.1016/j.fertnstert.2015.05.037

14 Zhang, Z. et al. Sperm retrieval outcomes of contralateral testis in men with nonobstructive azoospermia and unsuccessful unilateral microdissection testicular sperm extraction. Fertil Steril 121, 540–542 (2024). 10.1016/j.fertnstert.2023.11.023

15 Kim, W. J. et al. Intratesticular Peptidyl Prolyl Isomerase 1 Protein Delivery Using Cationic Lipid-Coated Fibroin Nanoparticle Complexes Rescues Male Infertility in Mice. ACS Nano 14, 13217–13231 (2020). 10.1021/acsnano.0c04936

16 Kormann, M. S. et al. Expression of therapeutic proteins after delivery of chemically modified mRNA in mice. Nat Biotechnol 29, 154–157 (2011). 10.1038/nbt.1733

17 Zhao, F. et al. Engineered nanoparticles potentials in male reproduction. Andrology (2024). 10.1111/andr.13729

18 Sahin, U., Kariko, K. & Tureci, O. mRNA-based therapeutics--developing a new class of drugs. Nat Rev Drug Discov 13, 759–780 (2014). 10.1038/nrd4278

19 Parhiz, H., Atochina-Vasserman, E. N. & Weissman, D. mRNA-based therapeutics: looking beyond COVID-19 vaccines. Lancet 403, 1192–1204 (2024). 10.1016/S0140-6736(23)02444-3

20 Vilpreux, C. et al. Sperm fertility in mice with oligo-astheno-teratozoospermia restored by in vivo injection and electroporation of naked mRNA. BioRxiv (2024). 10.1101/2023.12.12.571239

21 Wang, D., Tai, P. W. L. & Gao, G. Adeno-associated virus vector as a platform for gene therapy delivery. Nat Rev Drug Discov 18, 358–378 (2019). 10.1038/s41573-019-0012-9

22 Pupo, A. et al. AAV vectors: The Rubik’s cube of human gene therapy. Mol Ther 30, 3515–3541 (2022). 10.1016/j.ymthe.2022.09.015

23 Greig, J. A. et al. Integrated vector genomes may contribute to long-term expression in primate liver after AAV administration. Nat Biotechnol 42, 1232–1242 (2024). 10.1038/s41587-023-01974-7

24 Martins, K. M. et al. Prevalent and Disseminated Recombinant and Wild-Type Adeno-Associated Virus Integration in Macaques and Humans. Hum Gene Ther 34, 1081–1094 (2023). 10.1089/hum.2023.134

25 Zhang, N. N. et al. A Thermostable mRNA Vaccine against COVID-19. Cell 182, 1271–1283 e1216 (2020). 10.1016/j.cell.2020.07.024

26 Swingle, K. L. et al. Placenta-tropic VEGF mRNA lipid nanoparticles ameliorate murine pre-eclampsia. Nature 637, 412–421 (2025). 10.1038/s41586-024-08291-2

27 Han, E. L. et al. Peptide-Functionalized Lipid Nanoparticles for Targeted Systemic mRNA Delivery to the Brain. Nano Lett 25, 800–810 (2025). 10.1021/acs.nanolett.4c05186

28 Qiu, M. et al. Lung-selective mRNA delivery of synthetic lipid nanoparticles for the treatment of pulmonary lymphangioleiomyomatosis. Proc Natl Acad Sci U S A 119 (2022). 10.1073/pnas.2116271119

29 Du, S. et al. Cholesterol-Amino-Phosphate (CAP) Derived Lipid Nanoparticles for Delivery of Self-Amplifying RNA and Restoration of Spermatogenesis in Infertile Mice. Adv Sci (Weinh) 10, e2300188 (2023). 10.1002/advs.202300188

30 Wu, X. L. et al. The testis-specific gene 1700102P08Rik is essential for male fertility. Mol Reprod Dev 87, 231–240 (2020). 10.1002/mrd.23314

31 Struijk, R. B. et al. Simultaneous Purification of Round and Elongated Spermatids from Testis Tissue Using a FACS-Based DNA Ploidy Assay. Cytometry A 95, 309–313 (2019). 10.1002/cyto.a.23698

32 Kimura, Y. & Yanagimachi, R. Intracytoplasmic sperm injection in the mouse. Biol Reprod 52, 709–720 (1995). 10.1095/biolreprod52.4.709

33 Castrillon, D. H., Quade, B. J., Wang, T. Y., Quigley, C. & Crum, C. P. The human VASA gene is specifically expressed in the germ cell lineage. Proc Natl Acad Sci U S A 97, 9585–9590 (2000). 10.1073/pnas.160274797

34 Toyooka, Y. et al. Expression and intracellular localization of mouse Vasa-homologue protein during germ cell development. Mech Dev 93, 139–149 (2000). 10.1016/s0925-4773(00)00283-5

35 Lynn, A., Soucek, R. & Borner, G. V. ZMM proteins during meiosis: crossover artists at work. Chromosome Res 15, 591–605 (2007). 10.1007/s10577-007-1150-1

36 Baker, S. M. et al. Involvement of mouse Mlh1 in DNA mismatch repair and meiotic crossing over. Nat Genet 13, 336–342 (1996). 10.1038/ng0796-336

37 Wu, Y. et al. Correction of a genetic disease by CRISPR-Cas9-mediated gene editing in mouse spermatogonial stem cells. Cell Res 25, 67–79 (2015). 10.1038/cr.2014.160

38 Wang, Y. H. et al. Rescue of male infertility through correcting a genetic mutation causing meiotic arrest in spermatogonial stem cells. Asian J Androl 23, 590–599 (2021). 10.4103/aja.aja_97_20

39 Li, X., Sun, T., Wang, X., Tang, J. & Liu, Y. Restore natural fertility of Kit(w)/Kit(wv) mouse with nonobstructive azoospermia through gene editing on SSCs mediated by CRISPR-Cas9. Stem Cell Res Ther 10, 271 (2019). 10.1186/s13287-019-1386-7

40. Richter, M. F., et al. Phage-assisted evolution of an adenine base editor with improved Cas domain compatibility and activity. Nat Biotechnol 38, 883-891 (2020). 10.1038/s41587-020-0453-z

41 Pacesa, M. et al. Structural basis for Cas9 off-target activity. Cell 185, 4067–4081 e4021 (2022). 10.1016/j.cell.2022.09.026

42 Kim, D., Luk, K., Wolfe, S. A. & Kim, J. S. Evaluating and Enhancing Target Specificity of Gene-Editing Nucleases and Deaminases. Annu Rev Biochem 88, 191–220 (2019). 10.1146/annurev-biochem-013118-111730

43 Yuan, Y. et al. In vitro testicular organogenesis from human fetal gonads produces fertilization-competent spermatids. Cell Res 30, 244–255 (2020). 10.1038/s41422-020-0283-z

44 Medrano, J. V., Rombaut, C., Simon, C., Pellicer, A. & Goossens, E. Human spermatogonial stem cells display limited proliferation in vitro under mouse spermatogonial stem cell culture conditions. Fertil Steril 106, 1539–1549 e1538 (2016). 10.1016/j.fertnstert.2016.07.1065

45 Watanabe, S. et al. In Vivo Genetic Manipulation of Spermatogonial Stem Cells and Their Microenvironment by Adeno-Associated Viruses. Stem Cell Reports 10, 1551–1564 (2018). 10.1016/j.stemcr.2018.03.005

46 Liu, M. et al. HSF5 Deficiency Causes Male Infertility Involving Spermatogenic Arrest at Meiotic Prophase I in Humans and Mice. Adv Sci (Weinh) 11, e2402412 (2024). 10.1002/advs.202402412

47 Hordeaux, J. et al. Adeno-Associated Virus-Induced Dorsal Root Ganglion Pathology. Hum Gene Ther 31, 808–818 (2020). 10.1089/hum.2020.167

48 Van Alstyne, M. et al. Gain of toxic function by long-term AAV9-mediated SMN overexpression in the sensorimotor circuit. Nat Neurosci 24, 930–940 (2021). 10.1038/s41593-021-00827-3

49 Erles, K. et al. DNA of adeno-associated virus (AAV) in testicular tissue and in abnormal semen samples. Hum Reprod 16, 2333–2337 (2001). 10.1093/humrep/16.11.2333

50 Munro, A. P. S. et al. Safety and immunogenicity of seven COVID-19 vaccines as a third dose (booster) following two doses of ChAdOx1 nCov-19 or BNT162b2 in the UK (COV-BOOST): a blinded, multicentre, randomised, controlled, phase 2 trial. Lancet 398, 2258–2276 (2021). 10.1016/S0140-6736(21)02717-3

51 Stuart, A. S. V. et al. Immunogenicity, safety, and reactogenicity of heterologous COVID-19 primary vaccination incorporating mRNA, viral-vector, and protein-adjuvant vaccines in the UK (Com-COV2): a single-blind, randomised, phase 2, non-inferiority trial. Lancet 399, 36–49 (2022). 10.1016/S0140-6736(21)02718-5

52 Wang, Z. Y. et al. Transcriptome and translatome co-evolution in mammals. Nature 588, 642–647 (2020). 10.1038/s41586-020-2899-z

