## supplemental table 1-3 for "Testicular mRNA-LNP Delivery: A Novel Therapy for Genetic Spermatogenic Disorders"

Table S1. Primers for genotyping

| Primers | Sequence |
| --- | --- |
| *Msh5*-GT-F | 5’-CCCAAGGGATGAAAAGCCAC-3’ |
| *Msh5*-GT-R | 5’-GATACAGGGAGAGTAATGCGGTCTC-3’ |
| *Maps*-GT-F | 5′‐CCCGTTGCTCCGTGCATTTA‐3′ |
| *Maps*-GT-R | 5′‐TCTGCCTCCCGAGTGCTGTT‐3′ |

Table S2. Characterization of ionizable lipids and lipid nanoparticles

| Pool | Lipids | Mw | LNP | Size(nm) | PDI |
| --- | --- | --- | --- | --- | --- |
| Pool 1 | Lipid-001 | 706.15 | LNP-01 | 74 | 0.07 |
|  | Lipid-002 | 720.18 | LNP-02 | 77 | 0.06 |
|  | Lipid-003 | 734.2 | LNP-03 | 123 | 0.04 |
|  | Lipid-004 | 804.34 | LNP-04 | 77 | 0.13 |
|  | Lipid-005 | 944.61 | LNP-05 | 72 | 0.08 |
|  | Lipid-006 | 818.32 | LNP-06 | 76 | 0.09 |
|  | Lipid-007 | 618.04 | LNP-07 | 90 | 0.16 |
|  | Lipid-008 | 702.21 | LNP-08 | 79 | 0.18 |
|  | Lipid-009 | 874.56 | LNP-09 | 87 | 0.15 |
|  | Lipid-010 | 688.14 | LNP-10 | 85 | 0.15 |
| Pool 2 | Lipid-011 | 720.18 | LNP-11 | 96 | 0.08 |
|  | Lipid-012 | 804.34 | LNP-12 | 75 | 0.18 |
|  | Lipid-013 | 860.45 | LNP-13 | 92 | 0.13 |
|  | Lipid-014 | 734.16 | LNP-14 | 84 | 0.14 |
|  | Lipid-015 | 702.21 | LNP-15 | 91 | 0.15 |
|  | Lipid-016 | 674.15 | LNP-16 | 87 | 0.10 |
|  | Lipid-017 | 702.21 | LNP-17 | 90 | 0.19 |
|  | Lipid-018 | 618.04 | LNP-18 | 75 | 0.17 |
|  | Lipid-019 | 589.99 | LNP-19 | 88 | 0.13 |
|  | Lipid-020 | 888.5 | LNP-20 | 91 | 0.15 |
| Pool 3 | Lipid-021 | 776.29 | LNP-21 | 93 | 0.09 |
|  | Lipid-022 | 706.11 | LNP-22 | 76 | 0.09 |
|  | Lipid-023 | 734.16 | LNP-23 | 76 | 0.08 |
|  | Lipid-024 | 660.13 | LNP-24 | 83 | 0.17 |
|  | Lipid-025 | 674.15 | LNP-25 | 111 | 0.06 |
|  | Lipid-026 | 604.02 | LNP-26 | 94 | 0.18 |
|  | Lipid-027 | 772.34 | LNP-27 | 79 | 0.15 |
|  | Lipid-028 | 604.02 | LNP-28 | 90 | 0.17 |
|  | Lipid-029 | 748.14 | LNP-29 | 81 | 0.08 |
|  | Lipid-030 | 748.23 | LNP-30 | 91 | 0.18 |

Table S3. List of primary antibodies

| Antibody | Company | Catalog Number | Host | Dilution |
| --- | --- | --- | --- | --- |
| SYCP3 | Abcam | Ab97672 | Mouse | IF: 1: 200 |
| SYCP3 | Abcam | Ab15093 | Rabbit | IF: 1: 200 |
| SYCP1 | Abcam | Ab15090 | Rabbit | IF: 1: 200 |
| DMC1 | homemade | - | Rabbit | IF: 1: 200 |
| MLH1 | BD Pharmingen | 551092 | Mouse | IF: 1: 50 |
| DDX4 | Abcam | Ab13840 | Rabbit | IF: 1: 1000 |
| VIMENTIN | Servicebio | GB11192 | Rabbit | IF: 1: 200 |
| UCHL1 | Bio-Rad | MCA4750 | Mouse | IF: 1: 500 |
| EGFP | Abcam | Ab13970 | Chicken | IF: 1: 1000 |
| GFP | Invitrogen | A11122 | Rabbit | IF: 1: 1000 |
| AKAP3 | ProteinTech | 13907-1-AP | Rabbit | IF: 1: 200 |
| TP1 | ProteinTech | 17178-1-AP | Rabbit | IF: 1: 200 |
| GFP | Youke | YKCP 0312-01 | Mouse | WB: 1:1500 |
| ACTB-HRP | ProteinTech | HRP-66009 | Mouse | WB: 1:1000 |
